# PI3K inhibitor-free differentiation and maturation of human iPSC-derived arterial- and venous-like endothelial cells

**DOI:** 10.64898/2026.01.10.698801

**Authors:** Oliwia N. Mruk, Ralitsa R. Madsen

**Affiliations:** MRC Protein Phosphorylation and Ubiquitylation Unit, School of Life Sciences, University of Dundee, Dundee, DD1 5EH, United Kingdom; Cancer Research UK Scotland Institute, Glasgow, G61 1BD, United Kingdom; School of Cancer Sciences, University of Glasgow, Glasgow G61 1BD, United Kingdom

**Author notes:** Technical contact (Oliwia N. Mruk).

## Abstract

Congenital vascular malformations are commonly caused by aberrant, genetic activation of class I phosphoinositide 3-kinase (PI3K) signalling. Advances in mechanistic understanding and therapeutic targeting of these disorders will be accelerated by high-fidelity, human disease models. Building on a previously optimised differentiation strategy, we present a validated workflow for PI3K inhibitor-free generation of arterial- and venous-like endothelial cells from human induced pluripotent stem cells (iPSCs) under defined, xeno-free conditions. We further report experimental analyses of endothelial maturation under flow, culture duration-dependent stability, and downstream molecular and phenotypic characterisation. By providing a reproducible human system for mechanistic and translational studies, this platform will enable disease-relevant modelling of PI3K-driven vascular malformations, including *PIK3CA*-related overgrowth spectrum (PROS) and PTEN hamartoma tumour syndrome (PHTS).

## Background and Rationale

Mosaic activating mutations in *PIK3CA*, the gene encoding the catalytic subunit of phosphoinositide 3-kinase alpha (PI3Kα), cause 25-30% of congenital venous malformations (VMs) (Castillo et al. 2019), the most common type of vascular malformation. Although exceedingly rare, strongly activating *PIK3CA* mutations have also been reported in arteriovenous malformations (AVMs) in the context of complex overgrowth syndromes collectively termed *PIK3CA*-related overgrowth spectrum (PROS) (Kurek et al. 2012; Sterba et al. 2023; Green et al. 2024). Notably, AVMs are more common in the related *PTEN* hamartoma tumour syndrome (PHTS), a group of disorders caused by germline, loss-of-function mutations in the tumour suppressor PTEN (Tan et al. 2007; Gurunathan et al. 2020; Dykman et al. 2024; Abdelilah-Seyfried and Ola 2024). The latter dephosphorylates the PI(3,4,5)P_3_ second messenger produced by PI3Kα; accordingly, PHTS and PROS share partially overlapping clinical findings.

Better understanding of these PI3K-driven vascular pathologies, including the evidence for apparent developmental lineage skewing, and identification of improved treatment options requires access to high-fidelity, human “disease-in-a-dish”-based preclinical model systems. Several independent workflows for differentiation of human pluripotent stem cells (hPSC) into either arterial-(ALECs) or venous-like endothelial cells (VLECs) already exist but vary substantially with respect to duration, choice of differentiation components, extracellular matrix and lineage-specific marker evaluation (Sriram et al. 2015; Ang et al. 2022; Pan et al. 2025; Ang et al. 2025). Amongst these, the recent protocol by Ang/Loh et al. offers one of the most time-efficient and extensively benchmarked solutions, leading to robust VLEC and ALEC generation within 4-5 days (Ang et al. 2022; Loh et al. 2025). Crucially, however, it features the use of GDC0941, a class I PI3K inhibitor, which inhibits all p110 catalytic isoforms at the specified concentration of 2.5 µM. Thus, despite its robustness, the existing workflow would not be compatible with the modelling of developmental, PI3K-driven vascular malformations.

Building on Ang/Loh et al.’s work, we now present a robust PI3K inhibitor-free differentiation workflow for iPSC-derived ALEC and VLEC differentiation. We achieve this through the combined removal of insulin and GDC0941 at steps where both were previously required. Furthermore, we replace the use of Matrigel/Geltrex with synthetic laminin-511 E8 fragment as a more physiologically-relevant extracellular matrix (Miyazaki et al. 2012; Nguyen et al. 2016; Song et al. 2017), thereby facilitating fully defined differentiation culture conditions. Lastly, we present extensively optimised post-differentiation culture conditions for robust ALEC and VLEC maturation, including critical parameters such as cell density and flow. Altogether, we provide a reproducible, human-specific endothelial differentiation platform that will be of interest to developmental biologists, bioengineers, as well as researchers seeking access to scalable models of vascular diseases driven by genetic PI3K pathway activation.

## Methods

### Materials and Equipment

The main induced pluripotent stem cell (iPSC) line used in this work is the commercially-available WTC11 male line (RRID: CVCL_Y803). A derivative of the above parental line was obtained from the Allen Cell Explorer collection, engineered to contain a mGFP-tagged (monoallelic) vascular endothelial (VE)-cadherin (Allen Institute for Cell Science 2025; Cell Line ID: 126, Coriell #AICS-0126-068; WTC-mEGFP-CDH5-cl68)

A complete list of antibodies and reagents, along with their sources, is provided in **Tables 1 and 2**, respectively. The list of equipment and consumables used is provided in **Table 3**. The CDM2 Base Medium recipe is provided in **Table 4**, and stage-specific differentiation media recipes are provided in **Table 5**. The recipe for magnetic-activated cell sorting (MACS) buffer is also provided in **Table 6**. All the necessary reagents and equipment required for MACS are listed in **Table 7**.

**Table 1.**
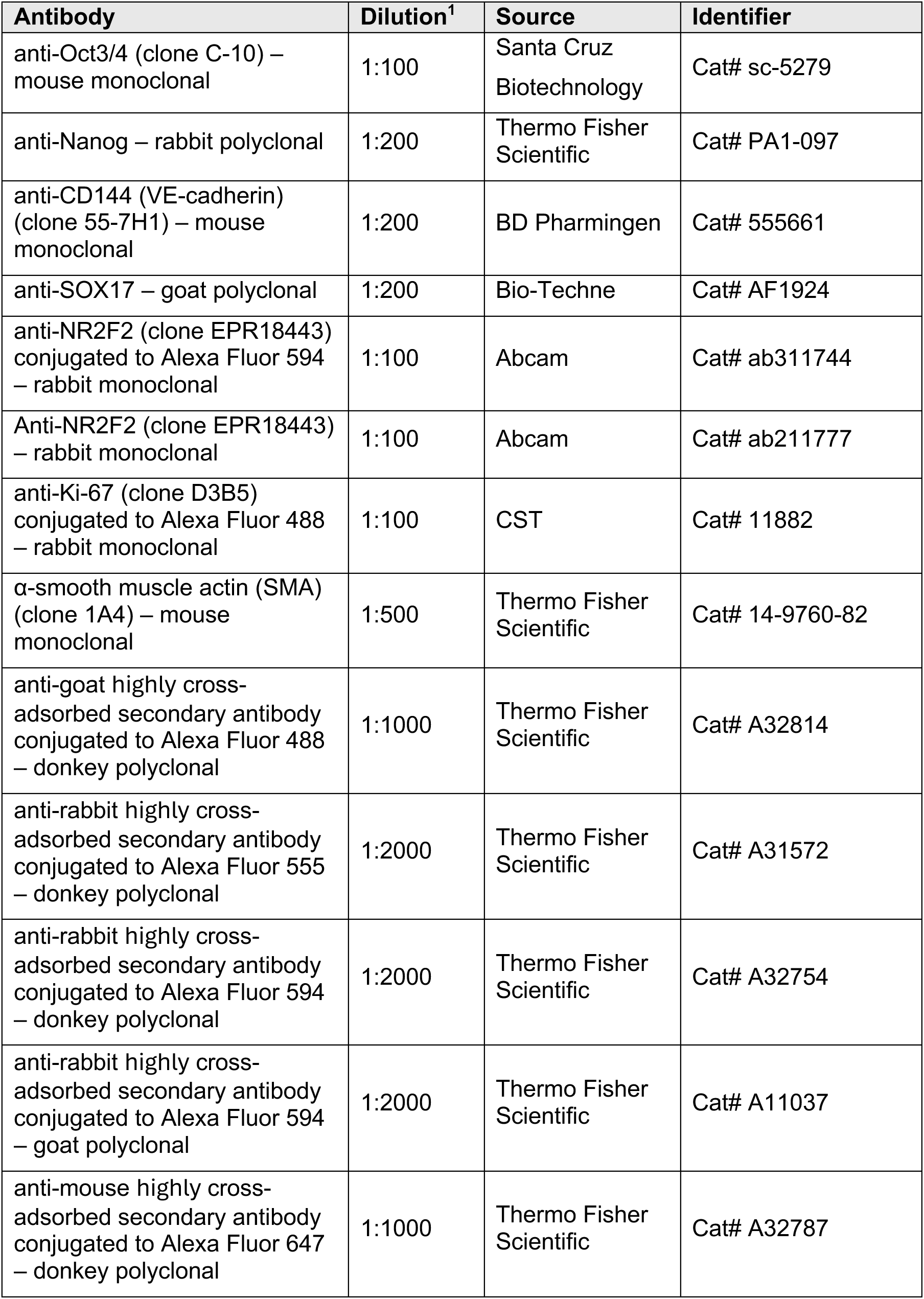
Primary and secondary antibodies. **^1^**All dilutions were prepared in 1X fish skin gelatin (FSG) in PBS/T (PBS + 0.05% Tween-20).

**Table 2.** Small molecules, proteins, chemicals and media.

| Reagent | Source | Identifier |
| --- | --- | --- |
| StemFlex Medium Kit | Thermo Fisher Scientific | Cat# A3349401 |
| ECMatrix-511 Silk E8 Laminin Substrate | Sigma-Aldrich | Cat# CC161-1050UG |
| Versene | Thermo Fisher Scientific | Cat# 15040066 |
| StemPro Accutase | Thermo Fisher Scientific | Cat# A1110501 |
| Dulbecco's phosphate-buffered saline (DPBS) | Thermo Fisher Scientific | Cat# 14190 |
| CTS PSC Cryomedium | Thermo Fisher Scientific | Cat# A4238801 |
| RevitaCell Supplement (10X) | Thermo Fisher Scientific | Cat# A2644501 |
| IMDM (1X) + GlutaMAX-I | Thermo Fisher Scientific | Cat# 31980030 |
| F12 (1X) + GlutaMAX-I | Thermo Fisher Scientific | Cat# 31765035 |
| Polyvinyl alcohol (PVA, 87-90% hydrolyzed, average molecular weight 30,000-70,000) | Sigma-Aldrich | Cat# P8136 |
| Chemically defined lipid concentrate | Thermo Fisher Scientific | Cat# 11905031 |
| Human transferrin | Sigma-Aldrich | Cat# T8158 |
| 1-Thioglycerol | Sigma-Aldrich | Cat# M6145 |
| Human recombinant insulin | Sigma-Aldrich | Cat# I9278 |
| Human recombinant BMP4 | Bio-Techne | Cat# 314-BPE-010 |
| Human recombinant BMP4 | Thermo Fisher Scientific | Cat# 120-05 |
| Human recombinant FGF-2 | Thermo Fisher Scientific | Cat# 100-18B |
| Human/mouse/rat recombinant Activin A | Thermo Fisher Scientific | Cat# 120-14P |
| CHIR99021 | Bio-Techne | Cat# 4423 |
| Human recombinant VEGF-165 | Thermo Fisher Scientific | Cat# 100-20 |
| NKH 477 | Sigma-Aldrich | Cat# N3290 |
| SB505124 | SelleckChem | Cat# S2186 |
| XAV939 | SelleckChem | Cat# S1180 |
| Ascorbic acid-2-phosphate | Sigma-Aldrich | Cat# A8960 |
| DMH1 | Sigma-Aldrich | Cat# 203646 |
| RO4929097 | SelleckChem | Cat# S1575 |
| PD0325901 | MRC PPU<br>Reagents &<br>Services, Dundee | - |
| PD0325901 | SelleckChem | Cat# S1036 |
| EGM-2 Endothelial Cell Growth Medium-2<br>BulletKit | Lonza | Cat# CC-3162 |
| Bovine Serum Albumin (BSA) 10 % in DPBS | Sigma-Aldrich | Cat# A1595 |
| Ethylenediaminetetraacetic (EDTA) acid<br>disodium salt solution | Sigma-Aldrich | Cat# 03690 |
| UltraPure 0.5M EDTA, pH 8.0 | Thermo Fisher<br>Scientific | Cat# 15575-038 |
| CD144 (VE-cadherin) MicroBeads, human | Miltenyi Biotec | Cat# 130-097-857 |
| Hoechst | Sigma-Aldrich | Cat#14533 |
| 10X PBS | Severn Biotech | Cat#20-7400 |
| Tween-20 | Promega | Cat#H5152 |
| 10X fish skin gelatin blocking agent (FSG) | Biotium | Cat#22010 |

**Table 3.** Equipment and consumables.

| Equipment/Consumable | Source | Identifier |
| --- | --- | --- |
| Nunc™ Cell-Culture Treated Multidishes – 6-well | Thermo Fisher Scientific | Cat# 140685 |
| Nunc™ Cell-Culture Treated Multidishes – 12-well | Thermo Fisher Scientific | Cat# 150628 |
| ViewPlate-96 TC black | PerkinElmer | Cat# 6005182 |
| PhenoPlate-96 TC black | Revvity | Cat# 6055300 |
| pluriStrainer Mini cell strainer, mesh size 40 µm | pluriSelect | Cat# 43-10040-50 |
| QuadroMACS Separator | Miltenyi Biotec | Cat# 130-090-976 |
| MACS MultiStand | Miltenyi Biotec | Cat# 130-042-303 |
| LS Columns | Miltenyi Biotec | Cat# 130-042-401 |
| RoboRocker 212 | NextAdvance | Cat# RR212-PL.G |

**Table 4.** CDM2 Base Medium stock and final concentrations of reagents. Store for up to 1 month at 4°C.

| Reagent | Solvent | Stock concentration | Final concentration |
| --- | --- | --- | --- |
| IMDM (1X) + GlutaMAX-I | n/a | 100% | 50% |
| F12 (1X) + GlutaMAX-I | n/a | 100% | 50% |
| Polyvinyl alcohol (PVA) | water | 50 mg/mL | 1 mg/mL |
| Chemically defined lipid concentrate | n/a | 100% | 1% (v/v) |
| Human transferrin | water | 30 mg/mL | 15 µg/mL |
| 1-thioglycerol | n/a | 11.5 M | 450 µM |

**Table 5.** Stage-specific differentiation media solutions and final concentrations of reagents. Store the solutions for up to 1 week at 4°C.

| Reagent | Solvent | Stock concentration | Final concentration |
| --- | --- | --- | --- |
| **Day 1: Mid primitive streak induction** |  |  |  |
| CDM2 Base Medium | n/a | n/a | n/a |
| Human recombinant insulin | n/a | 10 mg/mL | 0.7 µg/mL |
| Human recombinant BMP4 | 5 mM HCl | 20 µg/mL | 40 ng/mL |
| Human recombinant FGF-2 | 5 mM Tris-HCl | 30 µg/mL | 20 ng/mL |
| Human/mouse/rat recombinant Activin A | water | 100 µg/mL | 30 ng/mL |
| CHIR99021 | DMSO | 10 mM | 6 µM |
| **Day 2: Lateral plate mesoderm induction** |  |  |  |
| CDM2 Base Medium | n/a | n/a | n/a |
| Human recombinant BMP4 | 5 mM HCl | 20 µg/mL | 40 ng/mL |
| Human recombinant VEGF-165 | water | 100 µg/mL | 100 ng/mL |
| NKH 477 (forskolin analog) | water | 10 mM | 5 µM |
| SB505124 | DMSO | 10 mM | 2 µM |
| XAV939 | DMSO | 10 mM | 1 µM |
| Ascorbic acid-2-phosphate | water | 50 mg/mL | 200 µg/ml |
| **Day 3: Arterial induction** |  |  |  |
| CDM2 Base Medium | n/a | n/a | n/a |
| Human recombinant VEGF-165 | water | 100 µg/mL | 100 ng/mL |
| Human/mouse/rat recombinant Activin A | water | 100 µg/mL | 15 ng/mL |
| DMHI | DMSO | 1 mM | 250 nM |
| XAV939 | DMSO | 10 mM | 1 µM |
| Ascorbic acid-2-phosphate | water | 50 mg/mL | 200 µg/mL |
| **Day 3: Venous induction day 1** |  |  |  |
| CDM2 Base Medium | n/a | n/a | n/a |
| Human recombinant insulin | n/a | 10 mg/mL | 0.7 µg/mL |
| Human recombinant VEGF-165 | water | 100 µg/mL | 100 ng/mL |
| SB505124 | DMSO | 10 mM | 2 µM |
| DMH1 | DMSO | 1 mM | 250 nM |
| RO4929097 | DMSO | 10 mM | 2 µM |
| XAV939 | DMSO | 10 mM | 1 µM |
| Ascorbic acid-2-phosphate | water | 50 mg/mL | 200 µg/mL |
| **Day 4: Venous induction day 2** |  |  |  |
| CDM2 Base Medium | n/a | n/a | n/a |
| Human recombinant insulin | n/a | 10 mg/mL | 0.7 µg/mL |
| SB505124 | DMSO | 10 mM | 2 µM |
| RO4929097 | DMSO | 10 mM | 2 µM |
| CHIR99021 | DMSO | 10 mM | 1 µM |
| PD0325901 | DMSO | 1 mM | 500 nM |
| Ascorbic acid-2-phosphate | water | 50 mg/mL | 200 µg/mL |

**Table 6.**
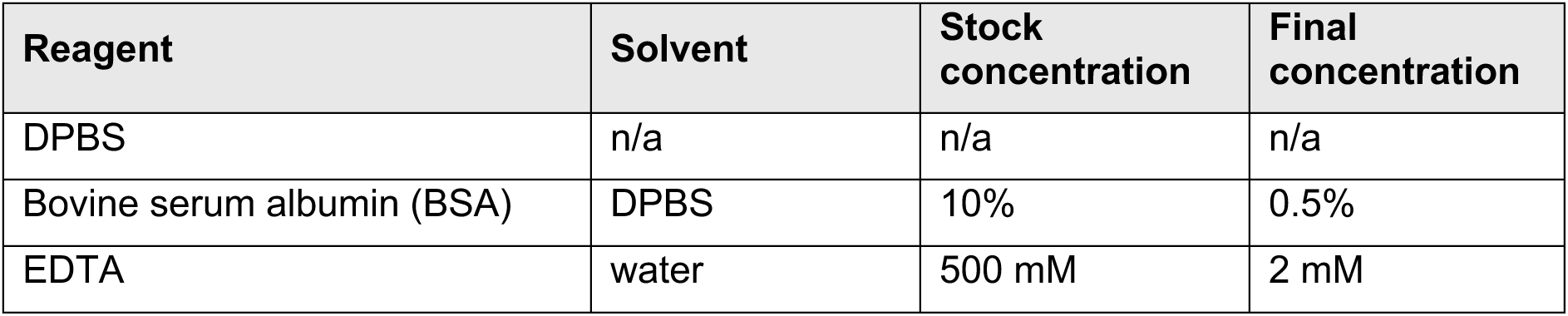
MACS buffer stock and final concentrations of reagents. Once sterile-filtered through an 0.2 µm PES membrane, store for up to 1 year at 4°C.

**Table 7.** Magnetic-activated cell sorting (MACS) reagents, in order of use in the workflow.

| Reagent/Equipment | Required for |
| --- | --- |
| DPBS | *Step 37* |
| StemPro Accutase | *Step 38* |
| MACS buffer | *Steps 39, 43, 47, 50, 52, 53, 54, 56* |
| CD144 (VE-cadherin) MicroBeads, human | *Steps 44, 45* |
| pluriStrainer Mini cell strainer(s), mesh size 40 µm | *Step 51* |
| LS column(s) | *Steps 52, 53, 54, 55* |
| QuadroMACS Separator | *Steps 52, 53, 54, 55* |
| 0.4% Trypan Blue | *Step 57* |
| EGM-2 Complete Medium | *Step 59* |

### General Experimental Preparation

#### Coating plates with ECMatrix-511 Silk E8 Laminin Substrate

**Timing:** Variable (1h – Overnight)

Induced pluripotent stem cells (iPSCs) and the differentiated cells resulting from this workflow are cultured on tissue culture-treated polystyrene plates (Nunclon Delta) coated with ECMatrix-511 Silk E8 Laminin Substrate. For this workflow, we describe initial culture and differentiation in a 6-well plate format, followed by expansion and maturation of pure endothelial cells in a 12-well plate format. The following procedure should be performed in a biosafety cabinet following appropriate aseptic technique.

1. Dilute the ECMatrix-511 Silk E8 Laminin Substrate stock from 0.5 mg/mL stock solution with sterile, ideally cold Dulbecco’s Phosphate-Buffered Saline (DPBS; no calcium, no magnesium) to achieve a 2.5 μg/mL working solution. Ensure immediate and sufficient mixing without vortexing.
2. Use 1 mL and 500 μL of the working solution to coat each well in a 6-well and 12-well plate, respectively. The final Laminin Substrate amount should correspond to 0.25 µg/cm^2^ and cover the well fully.

**Critical:** Gently move the plate in a “North-South-East-West” motion to ensure complete coverage of the bottom surface. If any areas are not covered (see **Figure 1B** for an example of uneven coating leading to uneven drying), laminin will not settle in those regions, and cells will fail to attach.

3. Incubate the plates at 37°C for 1h. Alternatively, leave the plates at room temperature for 3h or 4°C overnight (Parafilmed).

**Figure 1.**
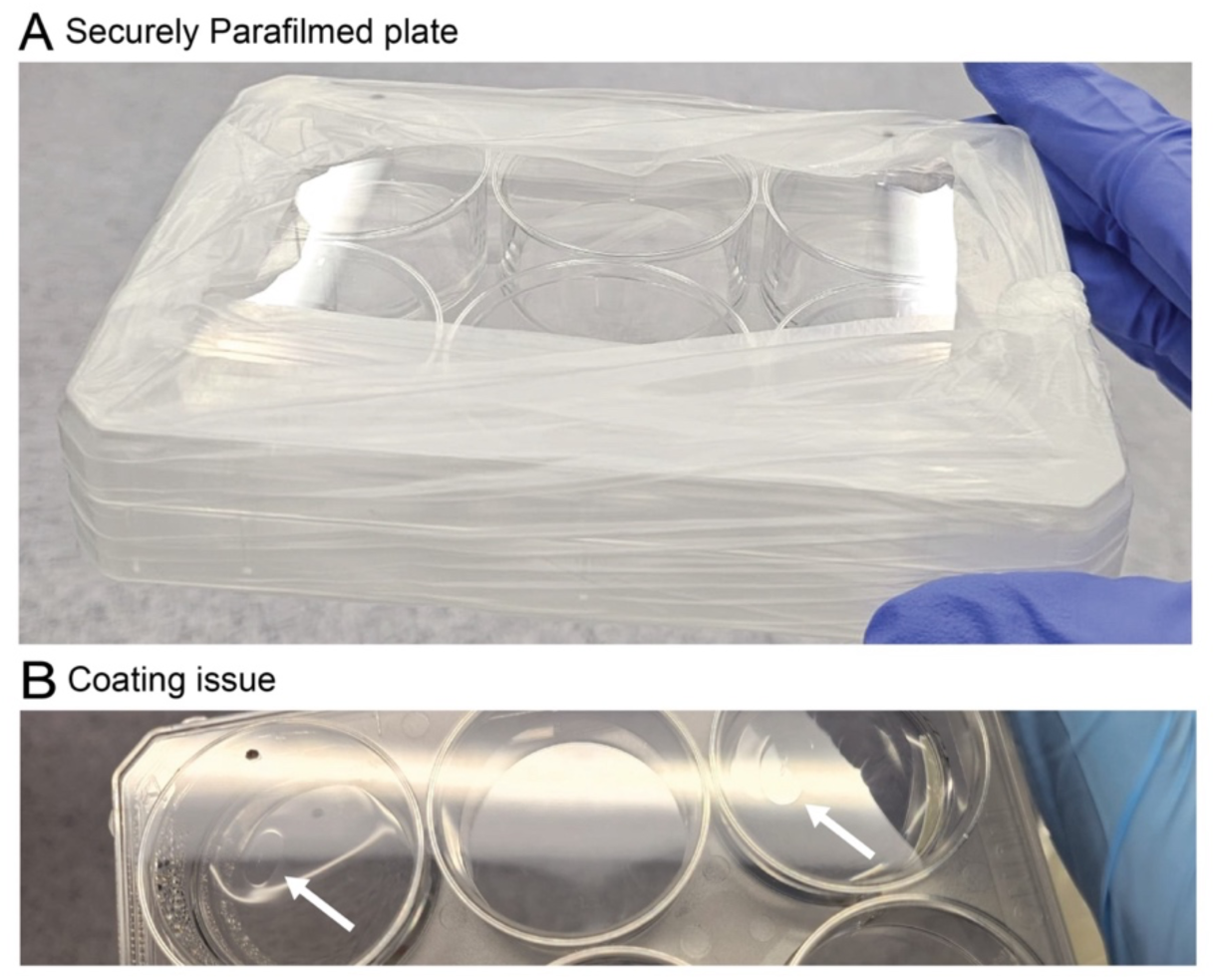
ECMatrix-511 Silk E8 Laminin Substrate coating and storage of plates. **A**. Securely Parafilmed plate, ready for longer-term storage at 4°C. **B**. Issue with coating; white arrows point to areas of the plate where the coating dried out and, therefore, cells will not attach.

**Optional**: Plates can be batch-coated in advance, secured with Parafilm and stored at 4°C for up to 14 days, provided the coating is even and has not evaporated across all or some of the well surface (**Figure 1**). Keep the plates on a level surface to avoid uneven coating. When ready to use, pre-warm the plates to room temperature for 30 minutes.

**Critical**: It is important to use the correct amount of laminin substrate per cm^2^. Do not leave ECMatrix-511 Silk E8 Laminin Substrate coated plates at 37°C for more than 24 h. This increases the risk of the coating drying out (**Figure 1B**), which will critically impact iPSC adherence, survival and differentiation. Topping up the coated wells with extra DPBS will have a similar effect due to dilution of the matrix and should be avoided.

#### Thawing iPSCs

**Timing:** 20 min

Here, we describe a routine iPSC thawing procedure for subsequent maintenance culture in StemFlex Medium and on ECMatrix-511 Silk E8 Laminin Substrate. StemFlex Medium refers to the StemFlex Basal Medium supplemented with the StemFlex Supplement at 1X, according to the manufacturer’s instructions. The following procedure should be performed in a biosafety cabinet following appropriate aseptic technique.

4. Prepare in advance:

a. The required volume of StemFlex Medium supplemented with RevitaCell Supplement diluted to 1X.

**Note**: RevitaCell Supplement (containing a Rho-associated, coiled-coil containing protein kinase (ROCK) inhibitor) prevents stress-induced apoptosis of hPSCs (Ohgushi et al. 2010) and is added to the medium to improve survival following thawing or when iPSCs are passaged as single cells. The medium volume needed will depend on the number of cells and their recovery efficiency. Generally, 10 mL of medium is prepared for immediate thawing before centrifugation, with an additional 4 mL required to fill two 6-wells with 2 mL of medium each. See **Figure 2** for details on layout and subsequent medium transfer. We routinely thaw c. 1 × 10⁶ cells/cryovial (clump-based) in this format, with excellent post-thawing recovery and an initial passaging within four days of thawing.

b. A 6-well plate with two wells coated with Laminin Substrate (as described in *Coating plates with ECMatrix-511 Silk E8 Laminin Substrate*). Once ready to use, gently aspirate the coating and add 2 mL of RevitaCell-supplemented StemFlex Medium per coated well. Place the plate in the humidified incubator to equilibrate to 37°C and the correct pH (5 % CO_2_ as required for bicarbonate-buffered media solutions).

**Figure 2.**
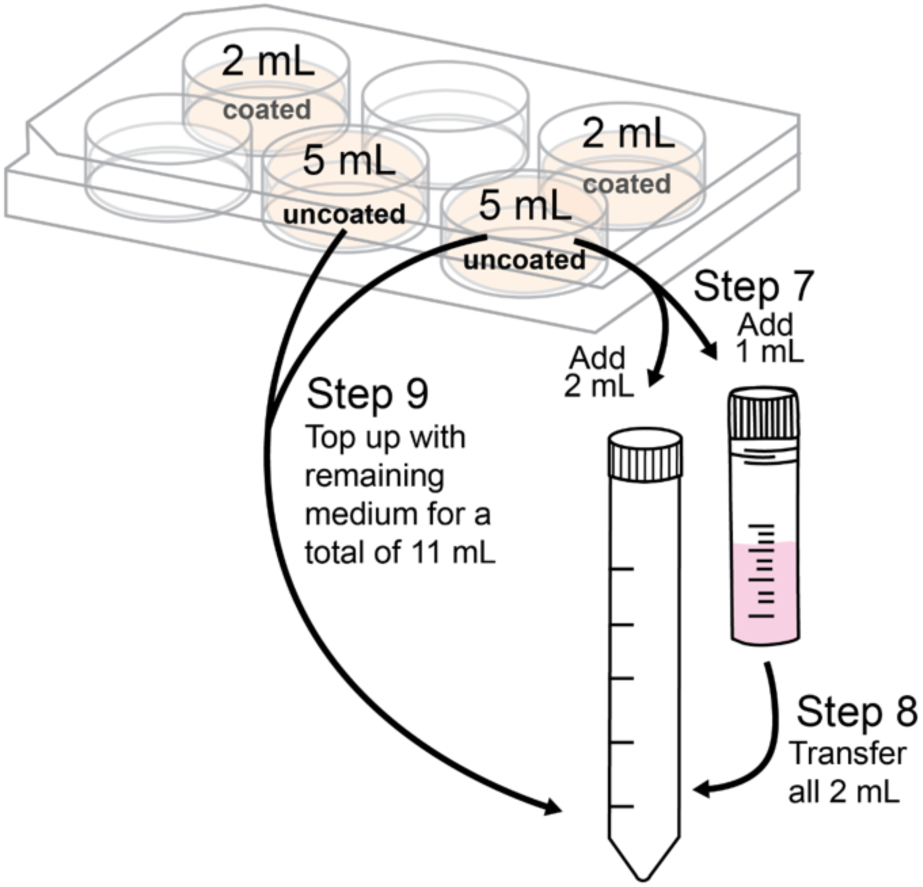
Schematic illustrating the procedure for *Steps 7–9* during medium and cell transfer for thawing.

**Note**: We recommend warming and pH-equilibrating the medium that will be used for the post-thaw wash of the cells. This can be done by adding a total of 10 mL of RevitaCell-supplemented StemFlex Medium across two of the non-coated 6-wells (see **Figure 2**) in the original plate to be equilibrated in the incubator.

c. A cryovial with 1 mL frozen cell suspension, kept on dry ice until ready to process.

**Note**: We do not recommend prolonged (> 1 week) storage of iPSCs at-80°C as this will compromise recovery post-thawing. The frozen stocks should be preserved in liquid nitrogen. Cryovials containing cells are stored in the liquid phase of liquid nitrogen; however, storage in the vapour phase is also suitable.

5. Take the cryovial, sterilise with 70% ethanol and transfer to thaw at 37°C in the humidified incubator until a tiny ice lump remains (approximately 4 minutes).

**Note**: We do not recommend the use of water baths for the thawing of antibiotic-free iPSC cultures. As an alternative to incubator thawing, hold the cryovial tightly in your hands until a small ice lump remains. We have never noticed an adverse effect on cell survival and use this approach with a wide range of cell lines.

6. Bring the cryovial to the biosafety cabinet together with the RevitaCell-supplemented StemFlex Medium.
7. Take up 3 mL of temperature and pH-equilibrated medium, then add 2 mL to the receiving 15 mL Falcon tube and, very slowly, the remaining 1 mL to the cryovial with cells.

**Critical**: This needs to be performed very slowly to avoid osmotic and thermal shock.

8. Immediately, and slowly, transfer the cell suspension to the 15 mL Falcon tube, just pre-filled with 2 mL of medium. Shake gently while dispensing to minimise osmotic and thermal shock.

**Optional**: Take up another 1-2 mL of medium from the prefilled wells to wash the cryovial to recover any cells remaining and combine with the cell suspension in the Falcon tube. Shake gently while dispensing.

9. Take up the remainder of the RevitaCell-supplemented StemFlex Medium from the uncoated 6-wells and gently add to the 15 mL Falcon tube with cells – you should now have 11 mL of cell suspension in total.
10. Return the 6-well plate with remaining medium in coated wells to the incubator to keep pH and temperature stable until ready to seed the cells.
11. Centrifuge the cell suspension at 200 g for 3 minutes in a swinging-bucket rotor centrifuge.
12. Remove the supernatant, leaving ∼50-100 μL of cell suspension behind and taking care not to aspirate the pellet.
13. Flick the tube gently to resuspend the pellet.
14. Take out the 6-well plate and use the medium in the coated well(s) to resuspend the pellet, taking care not to let the coating dry out. Resuspend the pellet gently and dispense it into the coated wells. Immediately after dispensing the cells, move the plate in a “North-South-East-West” direction to distribute the cells.

**Note**: We recommend taking 1 mL out of the 6-well; this will leave 1 mL in the well, not allowing the coating to dry out while the pellet is being resuspended. If unsure about the number of wells to use for cell thawing, start by seeding into one well, then inspect under the microscope. If the concentration of cells is too high or the clump size too large, take up the medium in the spare well, mix gently with the well containing cells and dispense 2 mL each across the two wells. The number of wells to use per thawing may need to be determined empirically based on the initial cell pellet size after centrifugation and knowledge of cell line-specific post-thaw recovery.

**Critical**: Take care to achieve a homogenous suspension of smaller cell clumps (3-5 cells) as opposed to single cells, as this helps the iPSCs survive. Avoid larger cell clumps of 20 cells or more, as this will increase the risk of differentiation or poor attachment.

15. Return the cells to the incubator and repeat the “North-South-East-West” movement for even dispersion.
16. Monitor cell recovery and remove RevitaCell Supplement after 24 h if colonies have formed successfully and appear to be growing.

**Note**: Our cryomedium (CTS PSC Cryomedium) does not contain FBS. If your cryomedium contains FBS, ensure to exchange the medium 4-8 h after thawing to remove any residual FBS, which may otherwise cause spurious differentiation. The cells should be re-fed with RevitaCell-supplemented medium up to 24 h post-thawing to aid recovery.

**Critical**: We recommend antibiotic-free iPSC culture to minimise any adverse effects of antibiotics on cell metabolism during growth. Extra care must therefore be taken to minimise the risk of contamination. This includes consistent use of filter tips, medium aliquots and immediate closing of tubes/plates after each use.

#### Daily iPSC maintenance

**Timing:** 10 min

It is important to assess the health of iPSC colonies continually and ensure no spontaneous differentiation is taking place. Healthy iPSCs are small and tightly packed into colonies. **Figure 3** provides examples of healthy iPSC colonies to use as a reference. As with all cell culture, the following procedure should be performed in a biosafety cabinet following appropriate aseptic technique.

17. Pre-warm StemFlex Medium to room temperature before starting.
18. Inspect the cells to assess their health and morphology.
19. Remove old medium from each 6-well containing cells.
20. Gently dispense 2 mL fresh StemFlex Medium per well. Dispense the medium along the side of the well to avoid direct disturbance of the cell monolayer.

**Figure 3.**
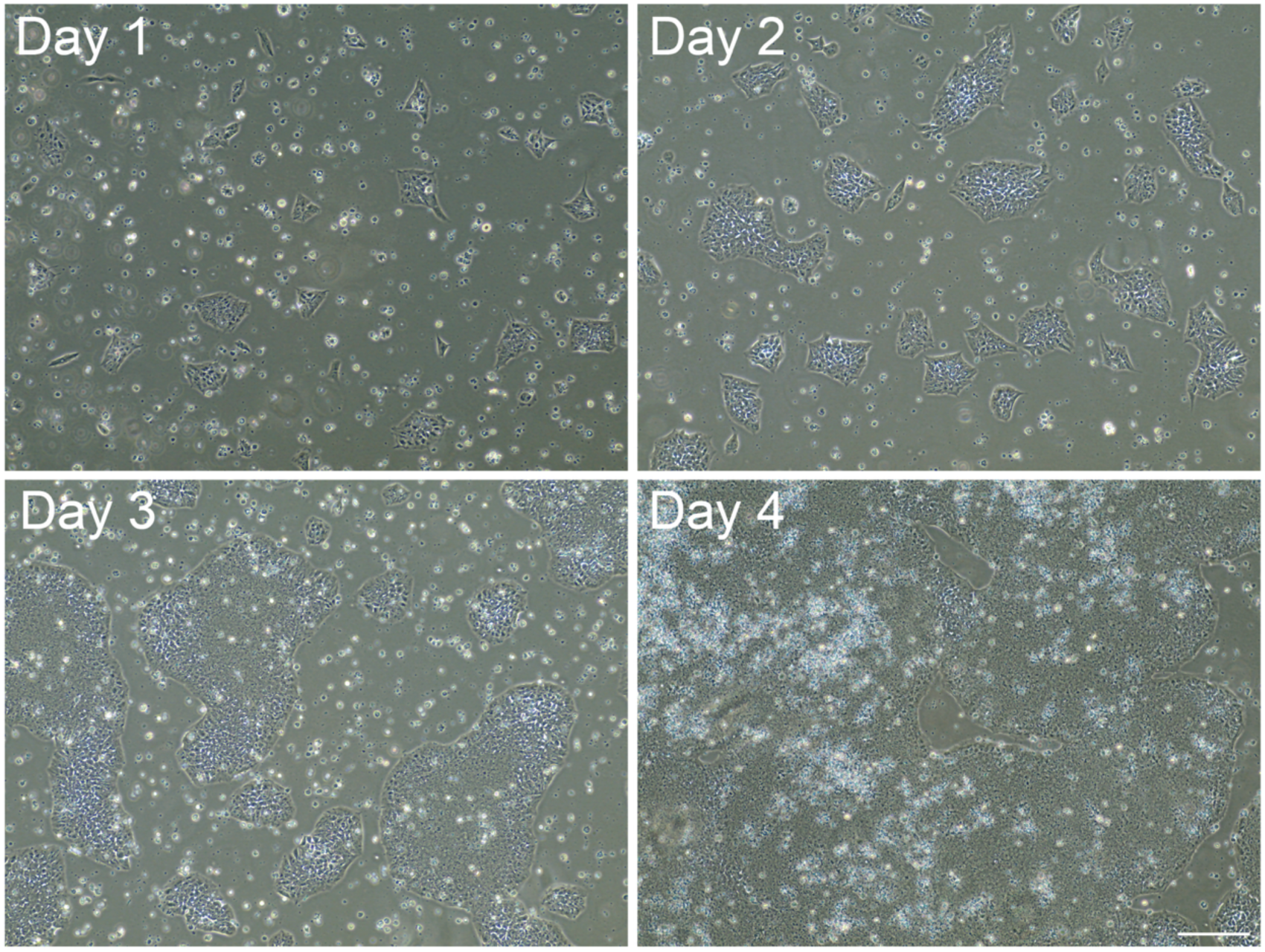
Representative micrographs illustrating healthy iPSC morphology over time. Cells were cultured in StemFlex on ECMatrix-511 Silk E8 Laminin Substrate. When colonies begin to merge (Day 4 example), the culture is ready for passaging. Scale bar: 200 μm.

**Note**: We typically achieve 100% undifferentiated iPSC cultures by ensuring consistent daily feeding times with freshly supplemented StemFlex Medium that has been aliquoted in advance, thereby avoiding opening/closing, pH buffering loss and repeated warming of a common stock bottle.

**Note**: When assessing cell confluency, it is advisable to also examine the edges of the well, as these often provide the most reliable indicator of overall readiness. Therefore, even if the centre of the well does not appear fully confluent, passaging is recommended if the edges have reached the appropriate confluency threshold.

#### Clump-based iPSC passaging with Versene (0.5 mM EDTA)

**Timing:** 15 min

When the iPSCs reach a confluency of ∼90% (**Figure 3**; Day 4), they require passaging. We perform clump-based passaging with Versene, which is gentle to the iPSCs, helps them survive and limits spontaneous differentiation. The following procedure should be performed in a biosafety cabinet following appropriate aseptic technique.

21. Prepare in advance:

a. A 6-well plate with the required number of wells coated with Laminin Substrate (as described in *Coating plates with ECMatrix-511 Silk E8 Laminin Substrate*). When ready to use, aspirate the coating and add 2 mL of StemFlex Medium per well. Return the plate to the incubator to allow equilibration to 37°C and the correct pH.
b. StemFlex Medium and Versene pre-warmed to room temperature before starting.

**Note**: We maintain two to three wells per cell line using different densities to achieve at least one optimal cell density for subsequent passaging, depending on the elapsed time (3-5 days). We recommend passaging iPSCs at least twice after thawing before using them for differentiation experiments.

22. Wash the 6-well to be split with 2 mL of DPBS.
23. Treat with 500 µL Versene and return to the incubator (37°C) for 6-8 minutes. Cells are considered sufficiently detached when they appear as shown in **Figure 4B**.

**Figure 4.**
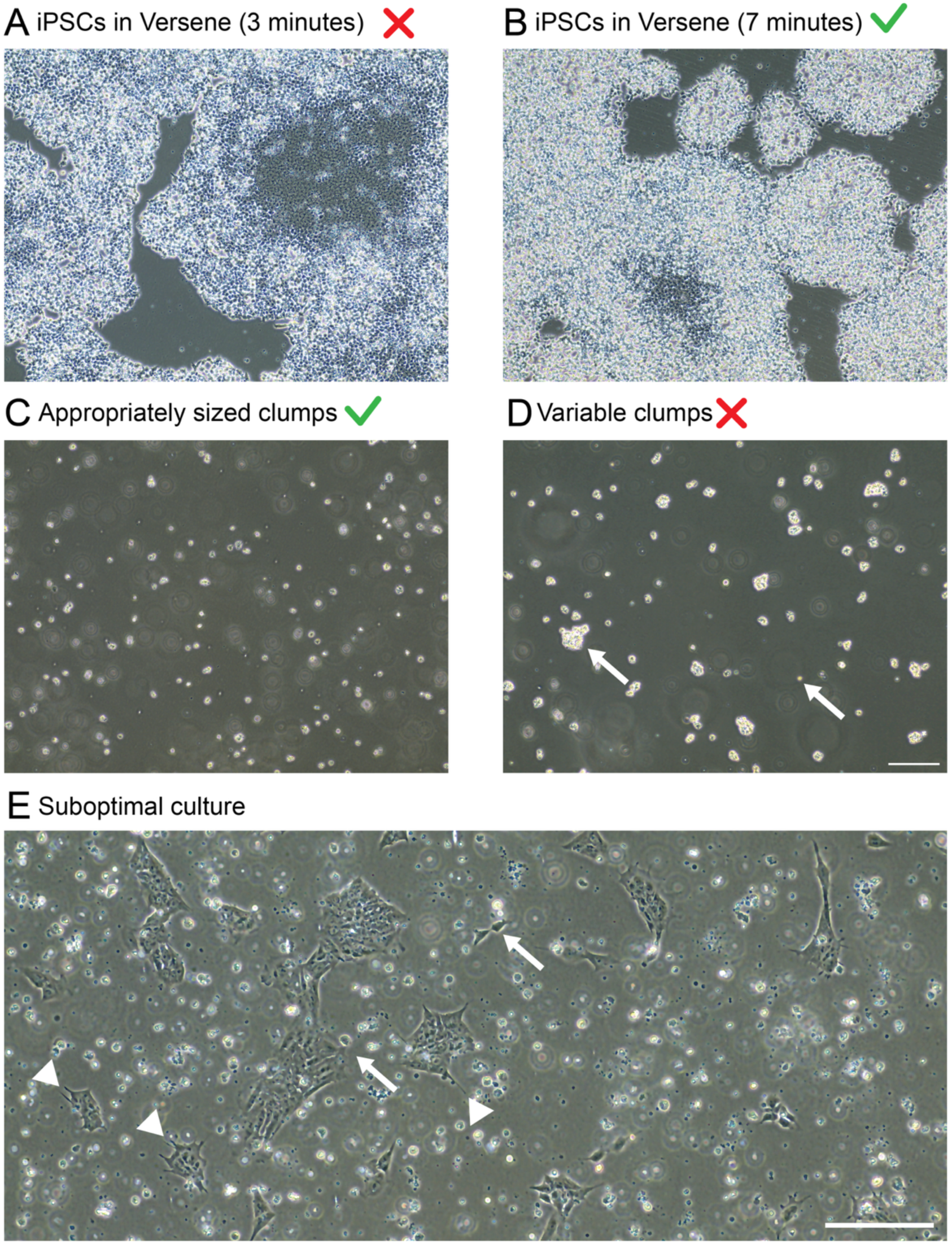
Clump-pased iPSC passaging. Representative micrographs illustrating optimal versus suboptimal results. White arrows indicate variability in clump size, which can lead to uneven cultures and increased susceptibility to differentiation. White arrowheads indicate projections that we use as markers of cell stress. Scale bar: 200 μm.

**Note**: Check cells under the microscope after the first 6 minutes to confirm the loosening of colonies. If the core of the colonies remains intact, return the plate to the incubator for another 1-2 minutes before checking again. Note that the rate of dissociation is slower the larger the colonies are, and will also vary as a function of iPSC line. Some iPSC lines may need longer than 8 minutes to dissociate fully.

**Critical**: Do not leave the iPSCs in Versene too long. The colonies should appear refractile with disrupted cell-cell junctions, but should not be lifting off by themselves. Additionally, in large confluent colonies, the centre may not detach fully (e.g. **Figure 4B**); if the majority of cells have loosened up, proceed with passaging. The strongly attached cells will remain behind, which helps limit spontaneous differentiation.

24. Return the plate with dissociated colonies to the biosafety cabinet, then open and tilt the plate at a 45° angle. Using a P1000 pipette, gently take up the Versene solution with cells, then dispense back across the well in a systematic motion that goes from one side to the other, from the top to the bottom of the well, whilst taking care not to pipette over the same area excessively. After doing this 4-5 times, you should observe that most cells have dissociated from the well, and the entire solution can now be collected into a sterile 1.5 mL Eppendorf.

**Critical**: Do not scratch the surface of the plate. Instead, wash the well gently just above to avoid agitating the cells.

25. Split according to your desired ratio by taking the required volume from the dissociated cell suspension into the well pre-filled with 2 mL of StemFlex Medium.

**Note**: For example, for a 1:20 split, you would take 25 µl of dissociated cells (out of 500 µL total volume). The usual ratios we use range between 1:5 and 1:25 without RevitaCell/ROCK inhibitor supplementation when cells are cultured on the recommended laminin substrate. Optimal ratios need to be determined empirically for each iPSC line. For the WTC11 iPSC line, an optimal split ratio of 1:20-1:25 will result in iPSC colonies that are ready for split 4-5 days later.

26. Immediately after dispensing the cells, gently move the plate according to a “North-South-East-West” movement for an even distribution.
27. Briefly inspect the cells under the microscope to confirm the correct clump size has been achieved (see example in **Figure 4C**).

**Note**: If cell clumps are larger than desired, e.g. as shown in **Figure 4D**, there is an increased risk of uneven attachment (**Figure 4E**) and unwanted differentiation. We recommend taking the plate back into the biosafety cabinet and gently pipetting up-and-down in the well with a P1000 pipette to break up larger clumps. Do not pipette more than 5 times to minimise excessive stress on the cells, and take care not to disturb the laminin coating.

28. Return the plate with cells back to the incubator and repeat the “North-South-East-West” movement for even dispersion.

#### Differentiation media preparation

**Timing:** Variable

Acquire all media and reagents as specified in *Materials and Equipment*. The following procedures should be performed in a biosafety cabinet following appropriate aseptic technique, and reagents should be sterile-filtered where necessary. Refer to **Supplementary File 1** for a pre-formatted Excel template with embedded formulas for media-specific component concentrations and volume calculations.

29. Reconstitute all reagents, as needed, following the manufacturer’s instructions.
30. Prepare CDM2 Base Medium by combining all reagents as specified under final concentration in *Materials and Equipment,* **Table 4**. Sterile-filter the solution through a 0.22 μm PES membrane.

**Note**: The CDM2 Base Medium can be stored in the fridge (4°C) for up to 1 month, provided that repeated opening/closing is kept to a minimum, or the medium subaliquoted, to preserve the 1-thioglycerol.

31. Prepare all stage-specific differentiation media solutions as specified under final concentration *Materials and Equipment*, **Table 5**.

**Note**: Store the stage-specific differentiation media solutions in the fridge (4°C) and use within a week of preparation. While differentiation medium may be stored for up to 1 week, optimal performance is achieved when the medium is supplemented fresh or shortly before use to limit degradation of sensitive components. Heating the medium is not recommended; aliquots should be equilibrated to room temperature before use.

32. Following magnetic-activated cell sorting (MACS), culture endothelial cells in EBM-2 Endothelial Cell Growth Medium-2 supplemented with BulletKit SingleQuots (EGM-2 Complete Medium). Omit the addition of GA-1000 (Gentamicin and Amphotericin) for antibiotic-free culture. Prepare according to the manufacturer’s instructions.

**Critical**: Arterial-like endothelial cells (ALECs) are cultured in EGM-2 Complete Medium. Venous-like endothelial cells (VLECs) require supplementation of EGM-2 Complete Medium with RO4929097 and SB505124, both at a final concentration of 2 μM, as described previously (Ang et al. 2022).

**Note:** EGM-2 Complete Medium may be prepared in advance and stored at 4 °C for up to 1 month according to the manufacturer’s recommendations. However, because human fibroblast growth factor basic (bFGF or FGF2) is relatively unstable, we recommend aliquoting the basal medium and supplements separately, with supplements stored at −80 °C. Complete medium should then be prepared as required and stored for up to 2 weeks after preparation to reduce the risk of activity loss due to prolonged storage at 4 °C. Similar precautions apply to StemFlex Medium preparation.

### Differentiation Procedure

This workflow outlines the generation of arterial-like and venous-like endothelial cells from iPSCs. Step-by-step instructions are provided for initial seeding, directed differentiation, purification using magnetic-activated cell sorting (MACS) and subsequent maturation of endothelial cells on flow (**Figure 5**). The following procedures should be performed in a biosafety cabinet following appropriate aseptic technique.

**Figure 5.**
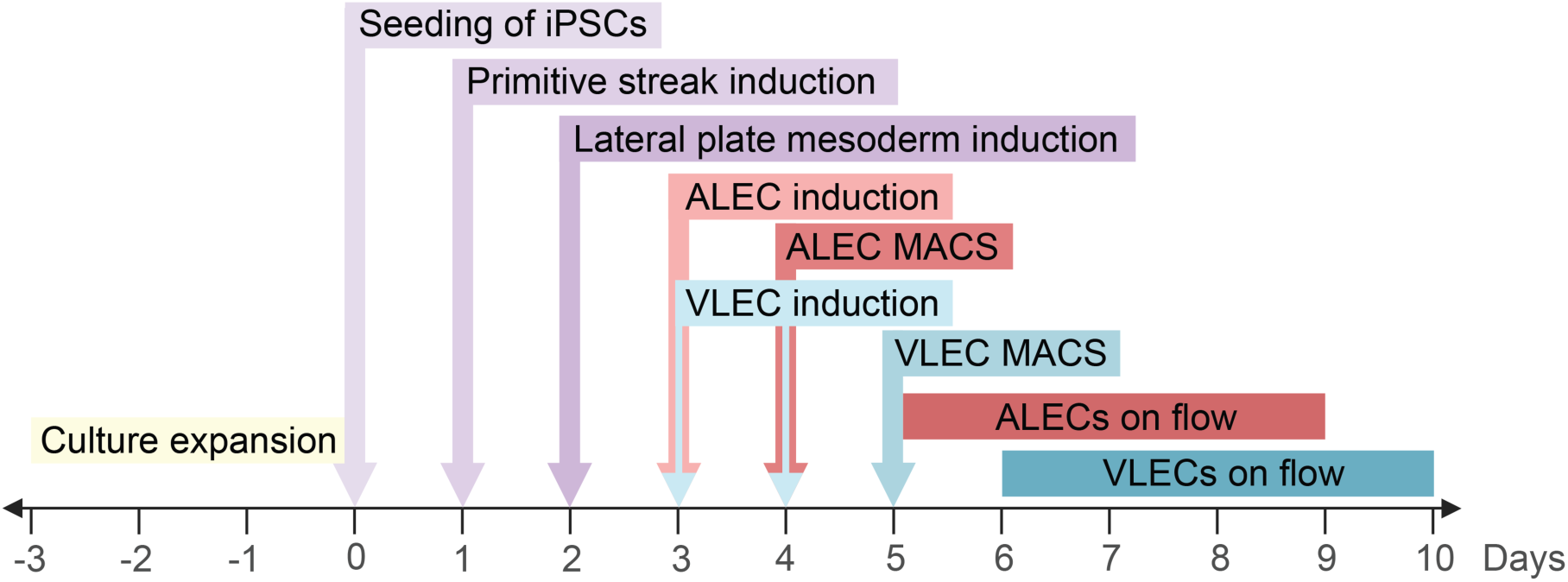
Differentiation timeline overview. ALEC, arterial-like endothelial cell; VLEC, venous-like endothelial cell.

**Critical**: No antibiotics are added to either iPSC or differentiating culture media. All steps must be performed under sterile conditions. If the medium appears cloudy or cells show unexpected signs of death, discard the culture and start with a fresh cell suspension. We also recommend weekly testing for mycoplasma.

#### Experimental seeding of iPSCs

**Timing:** 1 hour

This section outlines the initial seeding of iPSCs at optimal densities for growth and differentiation. The iPSCs should be harvested when subconfluent, as we have seen better differentiation results with cells that were around 70-80% dense.

1. Prepare in advance:

a. The required volume of StemFlex Medium supplemented with RevitaCell Supplement diluted to 1X, pre-warmed to room temperature.

**Note**: Each 6-well with iPSCs to be used for differentiation will be split with 500 µL of StemPro Accutase; to facilitate enzymatic inactivation, we collect the dissociated cells in StemFlex Medium with RevitaCell corresponding to 3 times the volume of Accutase. We therefore recommend aliquoting this collecting medium in advance. For example, if three iPSC 6-wells are to be split, prepare a tube containing 4.5 mL of medium. Additionally, each recipient well for differentiation should be seeded with a total of 2 mL of medium.

b. A tissue-culture treated 6-well plate, by coating the desired number of coated wells to receive cells (as described in *Coating plates with ECMatrix-511 Silk E8 Laminin Substrate*). Once the plates have been incubated for a sufficient period, gently aspirate the coating and add 1.5 mL of StemFlex Medium supplemented with RevitaCell per coated well. Put the plate in the incubator to equilibrate to 37°C and the correct pH.

**Note**: The number of wells to be seeded depends on the required cell numbers for downstream applications. At a minimum, we recommend seeding four to six 6-well plates. Although yields may vary, seeding from six 6-well plates typically allows for the generation of approximately six to twelve 12-wells for ALECs and four to eight 12-wells for VLECs.

c. A 1.5 mL Eppendorf pre-filled with 20 μL of 0.4% Trypan Blue for counting.
d. StemPro Accutase pre-warmed to room temperature before starting.

**Note**: We use StemPro Accutase for gentle single-cell-based passaging to allow for counting of cells for precise seeding.

2. Wash each 6-well to be split with 2 mL of DPBS.

**Note**: For seeding of six 6-wells (one full 6-well plate) for differentiation, start with three subconfluent 6-wells of iPSCs.

3. Treat each 6-well with 0.5 mL StemPro Accutase and return to the incubator (37°C) for 6-8 minutes.

**Note**: Check cells under the microscope after the first 6 minutes to confirm the loosening of colonies. If not ready, put back in the incubator for another minute before checking again. The detachment speed will be cell line specific.

4. Return the plate with dissociated colonies to the biosafety cabinet, then open and tilt the plate at a 45° angle. Using a P1000 pipette, gently take up the StemPro Accutase solution with cells, then dispense back across the well in a systematic motion that goes from one side to the other, from the top to the bottom of the well, whilst taking care not to pipette over the same area excessively. After doing this 4-5 times, you should observe that most cells have dissociated from the well, and the entire solution can now be collected into a tube filled with a sufficient amount of collecting medium, as outlined in *Step 1a* and the associated *Note*.
5. Take 20 µL of the cell suspension and add it to the Eppendorf tube containing Trypan Blue, outlined in *Step 1c*. Mix thoroughly, then load 10 µL into a haemocytometer. Count the cells manually.

**Note**: We recommend manual counting, as it gives more consistent seeding results compared with automated counters.

6. Calculate the volume of cells required to take forward for seeding each differentiation condition. For arterial differentiation, seed 45,000 cells/cm^2^; for venous differentiation, 30,000 cells/cm^2^. Refer to **Table 8** for the total number of cells required per well, including the 10% overage.

**Table 8.** Recommended cell numbers for differentiation initiation of arterial-like (ALECs) and venous-like endothelial cells (VLECs).

| Differentiation condition | Target cell seeding density (cells/cm<sup>2</sup>) | Target cells per 6-well (viable) | Cells per well with 10% overage | Cell suspension concentration (cells/mL) |
| --- | --- | --- | --- | --- |
| ALECs | 45,000 | 432,000 | 475,200 | 950,400 |
| VLECs | 30,000 | 288,000 | 316,800 | 633,600 |

**Note**: We add an additional 10% for the cell numbers to account for loss during centrifugation. We also calculate for seeding of an extra well, i.e. seven if we aim to seed six, to minimise the effect of pipetting loss following resuspension. For example, to seed cells into six wells in a 6-well plate format for ALEC differentiation, we take forward a total of 3.33 × 10^6^ cells for seven wells total, including the 10% overage.

7. Transfer the required volume of cells into a new 15 mL Falcon.
8. Centrifuge at 200 g for 3 minutes in a swinging-bucket rotor centrifuge.
9. Remove the supernatant, leaving ∼50-100 μL of cell suspension behind and taking care not to disturb the pellet.
10. Flick the tube gently to resuspend the pellet.
11. Add the required volume of StemFlex Medium supplemented with RevitaCell. For example, for seeding of six 6-wells, add a total of 3.5 mL (i.e. you will be seeding 500 µL into each well pre-filled with 1.5 mL medium, and as you have calculated for a total of seven wells, there should be about 500 µl left).
12. Mix gently to fully resuspend the cells in the medium.
13. Bring the recipient pre-filled and coated plate (see *Step 1b*) to the biosafety cabinet and dispense 500 μL of cell suspension per well. Immediately after dispensing the cells, move the plate in a “North-South-East-West” direction to distribute the cells.
14. Return the cells to the incubator and repeat the “North-South-East-West” movement for even dispersion.

#### Lineage-specific induction

**Timing:** 3 – 4 days (30 min per day)

This section outlines the steps for the directed differentiation from iPSCs into arterial-like (ALECs) and venous-like endothelial cells (VLECs) (**Figure 6**) using the CDM2 Base Medium with stage-specific supplementations (see also *Differentiation media preparation* and **Tables 4** and **5**). For reference, expected densities and morphologies of differentiating cells can be seen in **Figure 7**.

15. In advance, pre-warm the required volume of the relevant differentiation medium and DMEM/F-12 (no glutamine) to room temperature.

**Figure 6.**
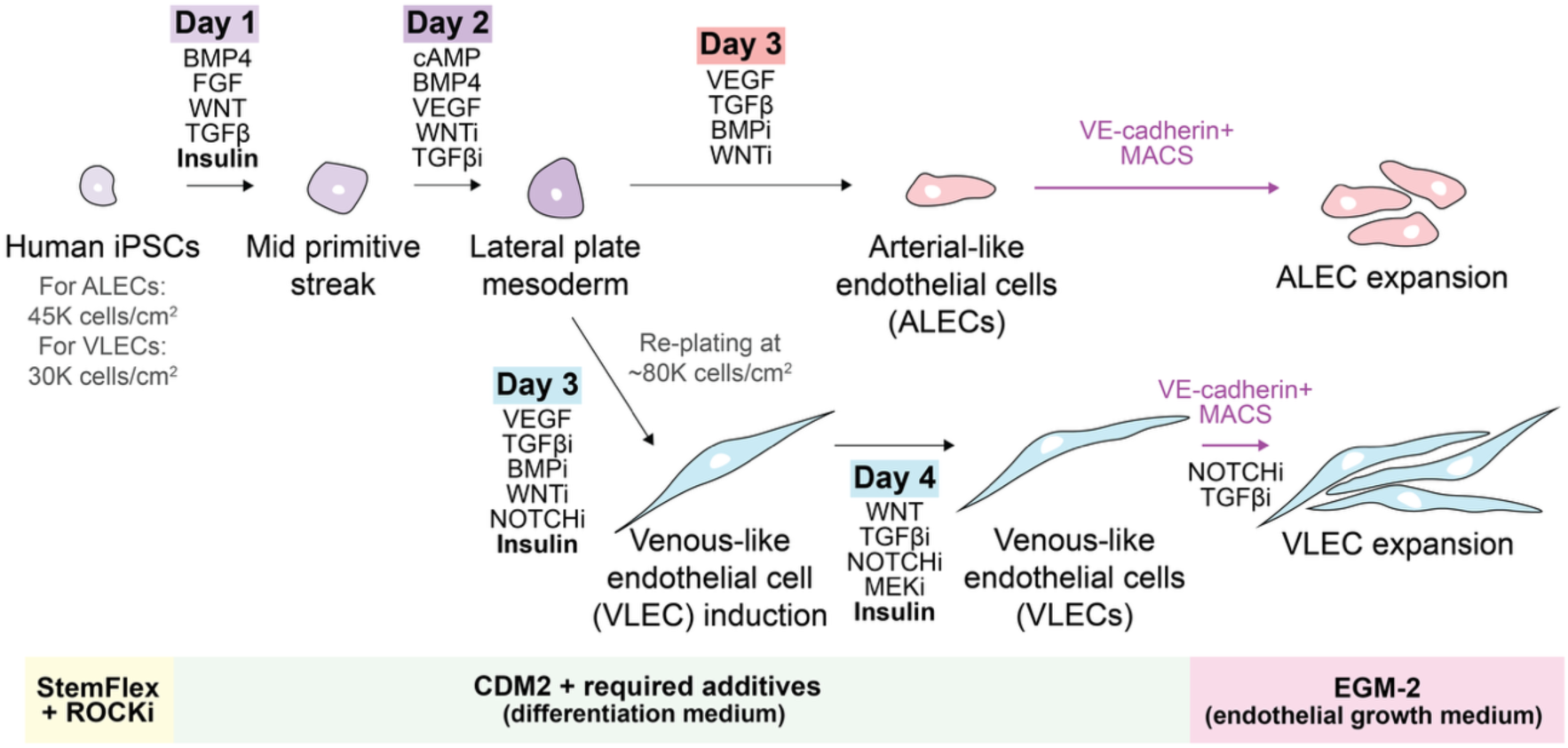
Overview of lineage-specific induction reagents and stages. Note the absence of insulin (physiological class IA PI3K activator) and GDC0941 (pharmacological class I PI3K inhibitor when used at 2.5 µM) at the lateral plate mesoderm and ALEC induction steps, a key difference from the original protocol by Ang et al. (2022).

**Figure 7.**
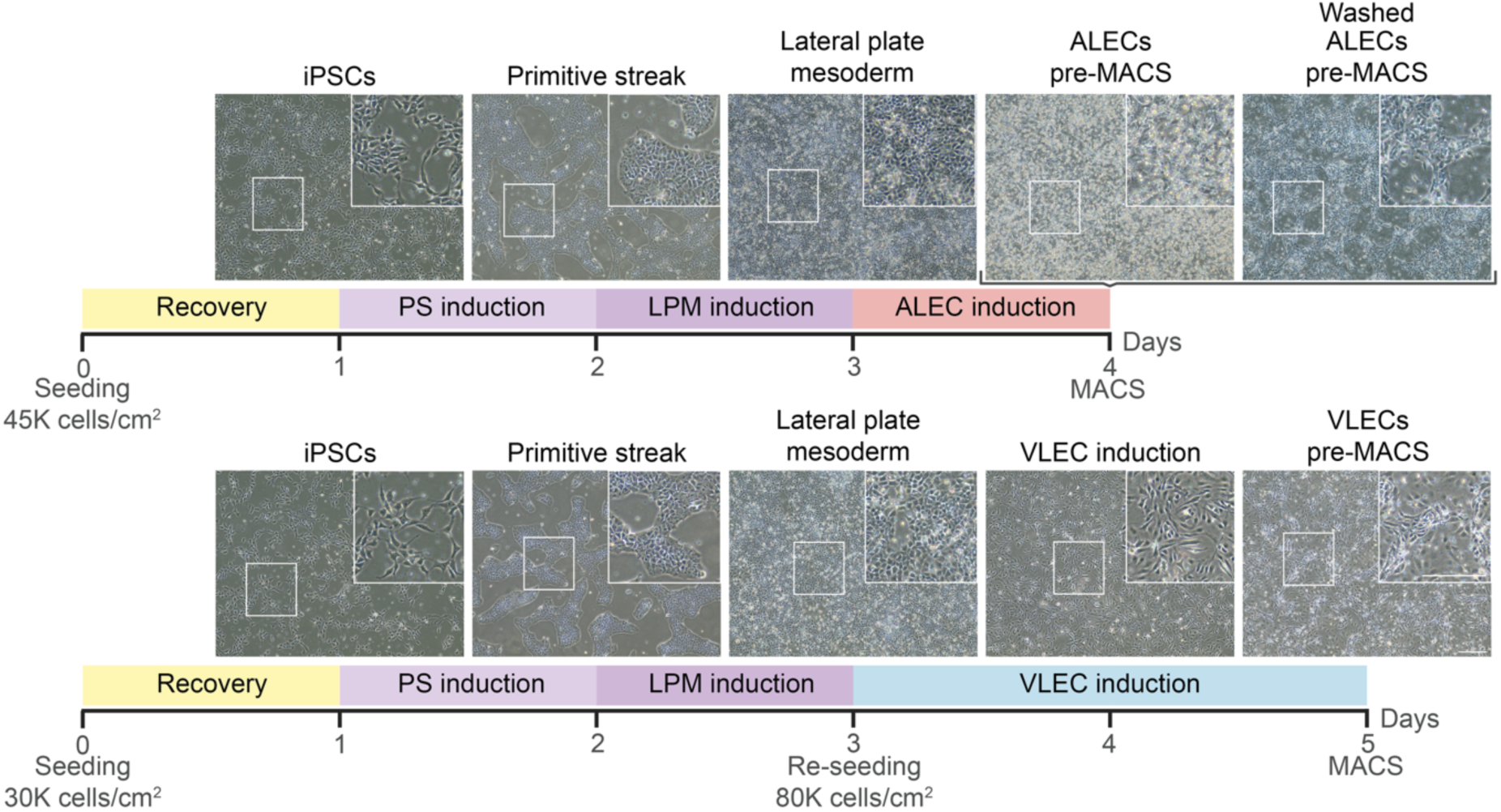
Expected cell densities and morphologies of cells at each differentiation stage. **Scale bar: 200 μm.**

**Critical**: Aliquot the media to maximise shelf life and maintain buffering capacity for consistent performance.

**Critical**: Individual medium changes at each differentiation step should occur at consistent 24 h intervals, with a maximum variation of ± 60 minutes allowed.

### Arterial lineage differentiation

16. Remove the previous day’s medium from each well and wash with 4 mL of DMEM/F-12 (no glutamine) per well.
17. Add 2 mL of appropriate differentiation medium per well.

**Critical**: When washing the cells, dispense 1 mL of medium first to each well and top up with the remaining 3 mL to avoid drying out and inconsistent timings across wells. Ensure you dispense the medium on the side of the well, quickly but gently, to avoid perturbing the cells. Cells should not be left in the wash medium for extended periods. Exposure time should be minimised as much as possible, as prolonged incubation can negatively impact the downstream differentiation performance.

**Note**: One day after arterial differentiation induction, a lot of cell death is typically observed along with the emergence of tube-like structures connecting individual cell clusters (see **Figure 7** for ALECs pre-MACS before and after a double wash with DPBS to remove debris).

### Venous lineage differentiation

18. For Day 1 (mid primitive streak) and Day 2 (lateral plate mesoderm) inductions: Remove the previous day’s medium from each well and wash with 4 mL of DMEM/F-12 (no glutamine) per well as described in *Lineage Specific Induction Steps 16 and 17*.
19. Add 2 mL of the appropriate differentiation medium per well.

**Critical**: As previously outlined, these steps should be repeated every 24 h until the iPSCs have committed to the desired lineage. It is very important to maintain this timing, with a maximum variation of ± 60 minutes allowed.

Following differentiation into lateral plate mesoderm, the cells require re-seeding to adjust for cell density and improve the venous differentiation efficiency, as described previously (Ang et al. 2022).

20. Prepare in advance:

a. The required volume of Day 3 (venous induction day 1) medium, pre-warmed to room temperature.

**Note**: Each 6-well will be split with 500 µL of StemPro Accutase – this must be inactivated by adding 3 times the volume of Day 3 medium to stop enzymatic activity. We therefore recommend aliquoting this collecting medium in advance. For example, if six wells are to be split, prepare a tube containing 9 mL of collecting medium. Additionally, each recipient well should be seeded with a total of 2 mL of medium.

b. A 6-well plate with the desired number of coated wells (as described in *Coating plates with ECMatrix-511 Silk E8 Laminin Substrate*). Gently aspirate the coating and add 1 mL of Day 3 (venous induction day 1) differentiation medium per coated well. Transfer the plate to the incubator to equilibrate to 37°C and the correct pH.

**Note**: The number of wells to be seeded into will depend on the number of wells pre-seeded and their yield following dissociation. We are typically able to re-seed cells into nine 6-wells when we collect cells from a full 6-well plate.

c. A 1.5 mL Eppendorf pre-filled with 20 μL of 0.4% Trypan Blue for counting.

21. Pre-warm StemPro Accutase to room temperature before starting.
22. Wash each 6-well to be split with 2 mL of DPBS.
23. Treat each well with 0.5 mL StemPro Accutase and return to the incubator (37°C) for around 5 minutes.

**Note**: Check cells under the microscope after the first 3-4 minutes to confirm the loosening of cells. The detachment of these cells is much faster than iPSCs, so prepare to process them quickly.

24. Return the plate with dissociated cells to the biosafety cabinet, then open and tilt the plate at a 45° angle. Using a P1000 pipette, gently take up the StemPro Accutase solution with cells and then dispense back across the well in a systematic motion that goes from one side to the other, from the top to the bottom of the well, whilst taking care not to pipette over the same area excessively. The cells dissociate easily and can now be collected into a tube filled with a sufficient amount of medium, as outlined in *Step 20a*.

**Critical**: Do not scratch the surface of the plate. Instead, wash the well gently just above to avoid agitating the cells.

25. Take 20 µL of the cell suspension and add to the Eppendorf tube containing Trypan Blue, as outlined in *Step 20c*. Mix thoroughly, then load 10 µL into a haemocytometer. Count the cells manually.
26. Calculate the volume of cells required for 80,000 cells/cm^2^ re-seeding. Refer to **Table 9**, including the 10% overage and volume for an extra well for seeding as described in *Step 6*.
27. Take up the required volume of cells and transfer to a new tube.

**Table 9.** Recommended cell numbers for re-seeding of differentiating venous-like endothelial cells (VLECs).

| Differentiation condition | Target cell seeding density (cells/cm<sup>2</sup>) | Target cells per 6-well (viable) | Cells per well with 10% overage | Cell suspension concentration (cells/mL) |
| --- | --- | --- | --- | --- |
| VLECs re-seeding | 80,000 | 768,000 | 844,800 | 844,800 |

**Note**: For example, to seed nine 6-wells, collect 8.03 × 10^6^ cells for nine and a half wells total (including the overage).

28. Centrifuge the required volume of cell suspension at 200 g for 3 minutes in a swinging-bucket rotor centrifuge.
29. Remove the supernatant, leaving ∼50-100 μL of cell suspension behind and taking care not to disturb the pellet.
30. Flick the tube gently to resuspend the pellet.
31. Add the required volume of Day 3 (venous induction day 1) medium. For example, for seeding of nine 6-wells (nine and a half wells with extra), add a total of 9.5 mL.
32. Mix gently to fully resuspend the cells in the medium.
33. Bring the recipient pre-filled plate with coated wells and pre-dispensed medium (see *Step 20b*) to the biosafety cabinet and dispense 1 mL of well-mixed cell suspension per well. Immediately after dispensing the cells, move the plate in a “North-South-East-West” direction to distribute the cells.
34. Return the cells to the incubator and repeat the “North-South-East-West” movement for even dispersion.
35. Following 24 h of incubation, remove the previous day’s medium from each well and wash with 4 mL of DMEM/F-12 (no glutamine) as described previously in *Lineage Specific Induction Steps 16 and 17,* then add 2 mL of Day 4 (venous induction day 2) medium per well.

#### Magnetic-activated cell sorting (MACS) for CD144 (VE-cadherin) to enrich for endothelial cell populations

**Timing:** 2 – 4 hours

This section provides step-by-step instructions for enrichment of pure endothelial cell populations using magnetic microbeads conjugated to an anti-VE-cadherin antibody.

36. Prepare in advance:

a. MACS buffer, kept cold in the fridge (4°C) or on ice during the duration of the procedure (see **Table 6** for recipe).
b. Add 900 µL of cold MACS buffer to a 1.5 mL Eppendorf tube for each well that will be processed. For example, if processing 6 wells per cell line, then prepare 6 Eppendorf tubes. Keep the tubes on a cold block.
c. Additional 1.5 mL Eppendorf tubes, pre-filled with 50 μL of cold MACS buffer and labelled with the sample name. Keep the tubes on a cold block.

**Note**: The number of tubes will depend on the number of processed wells. We have found that, for the best results, cells from a maximum of three 6-wells should be used per MACS column. For example, if processing six 6-wells per sample, prepare two tubes with 50 µl MACS buffer each.

d. Additional 1.5 mL Eppendorf tube(s) pre-filled with 20 μL of 0.4% Trypan Blue for counting.

**Optional**: Only one tube per clone is required; however, we also recommend preparing an additional tube per clone to count flow-through cells for calculations of differentiation efficiency.

e. EGM-2 Complete Medium to accept either arterial or venous-like endothelial cells, pre-warmed to room temperature, to collect ALECs and VLECs post-MACS.

**Critical**: As outlined in *Differentiation media preparation, Step 32 Critical*, the EGM-2 Complete Medium to accept VLECs requires the addition of RO4929097 and SB505124.

f. StemPro Accutase, pre-warmed to room temperature.
g. A laminin-coated tissue-culture treated 12-well plate(s) with laminin coating (as described in *Coating plates with ECMatrix-511 Silk E8 Laminin Substrate)* to receive cells post-MACS.

**Note**: The number of wells to be seeded into will depend on the efficiency of the differentiation process, the starting number of cells and the amount of cell loss during MACS. For ALECs, we recommend coating up to twelve wells in a 12-well plate if processing a total of six 6-wells for MACS; for VLECs, we recommend coating up to eight wells in a 12-well plate if processing a total of nine 6-wells for MACS post-reseeding. This may vary if using a different cell line and should be determined empirically.

h. All else required inside the biosafety cabinet for MACS, including specific materials, beads and magnets. Please refer to **Figure 8** and **Table 7** for setup guidance.

37. Wash each 6-well to be dissociated with 2 mL of DPBS.

**Figure 8.**
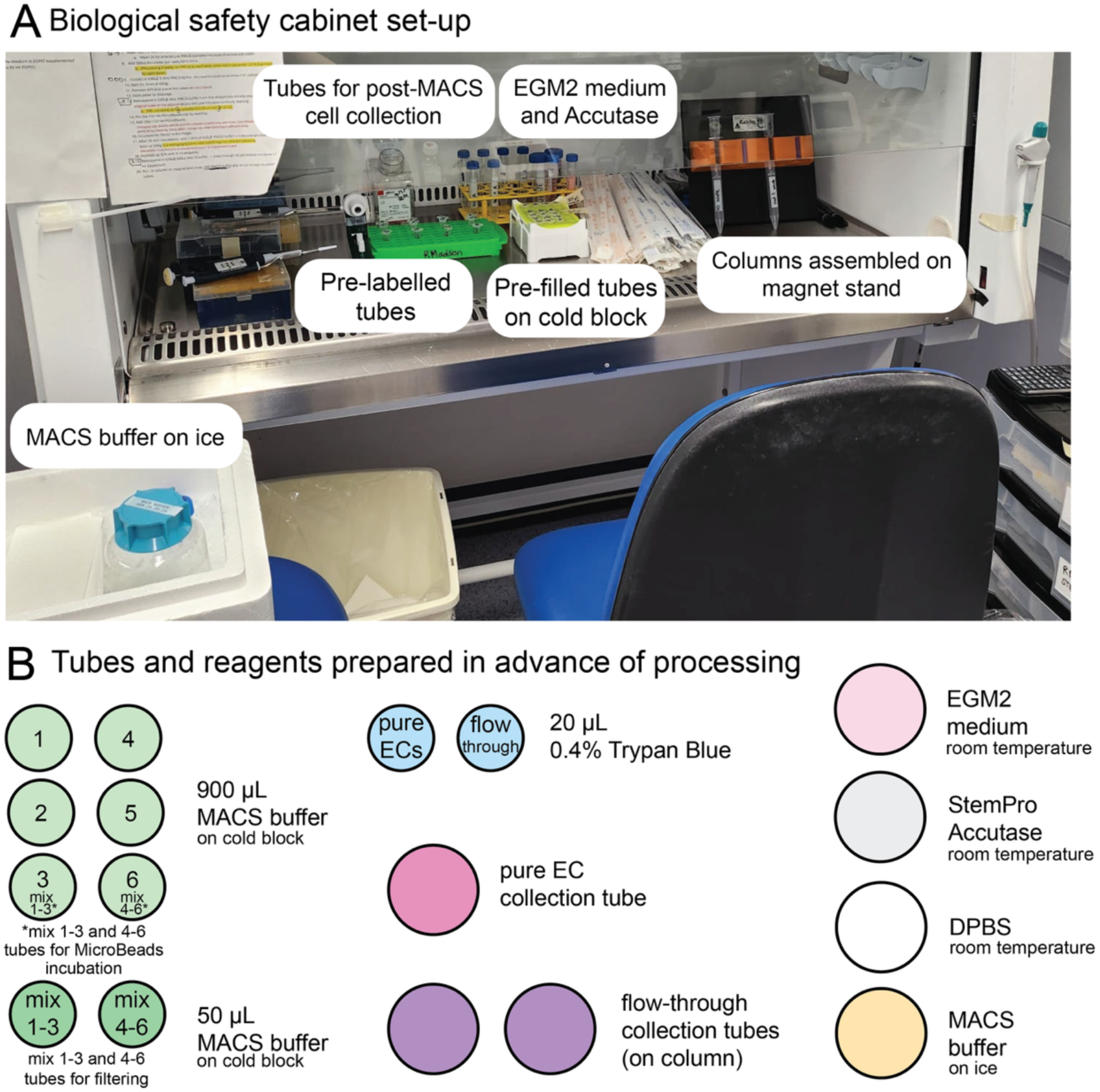
Set-up for MACS a biological safety cabinet. The set-up shown here is for processing of the same culture seeded across a full 6-well plate. Smaller-sized circles represent 1.5 mL Eppendorf tubes, and bigger-sized circles represent larger collection tubes, such as 15 mL or 50 mL Falcon tubes (depending on volumes needed). Abbreviations: EC: endothelial cell; MACS: magnetic activated cell sorting.

**Critical**: Due to more extensive cell death at the last step of arterial differentiation, it is important to wash the cells with DPBS twice.

38. Treat with 500 µL StemPro Accutase and return to the incubator (37°C) for 5 minutes.
39. Return the plate with dissociated colonies to the biosafety cabinet, then open and tilt the plate at a 45° angle. Using a P1000 pipette, gently take up the StemPro Accutase solution with cells, then dispense back across the well in a systematic motion that goes from one side to the other, from the top to the bottom of the well, whilst taking care not to pipette over the same area excessively. The cells dissociate easily, and each well can now be collected in the tube pre-filled with 900 µL of MACS buffer, stored on a cold block.

**Critical**: Do not scratch the surface of the plate. Instead, wash the well gently just above to avoid agitating the cells.

40. Centrifuge at 200g for 3 minutes in a swinging-bucket rotor centrifuge for efficient pelleting.

**Note**: If your swinging-bucket rotor centrifuge does not have adaptors for Eppendorfs, use empty 15 mL Falcon tubes as adaptors but do not exceed 200 g in centrifugation speed or your Eppendorf will fall through and get stuck in the 15 mL Falcon.

41. Remove the supernatant and keep the tubes on a cold block.
42. Flick the tubes gently to re-suspend the pellets.
43. Resuspend the pellets in the 50 μL of cold MACS buffer prepared earlier (*Step 36c*;

Figure 9). Do not discard the original 1.5 ml Eppendorf tube containing the 50 µl.

**Figure 9.**
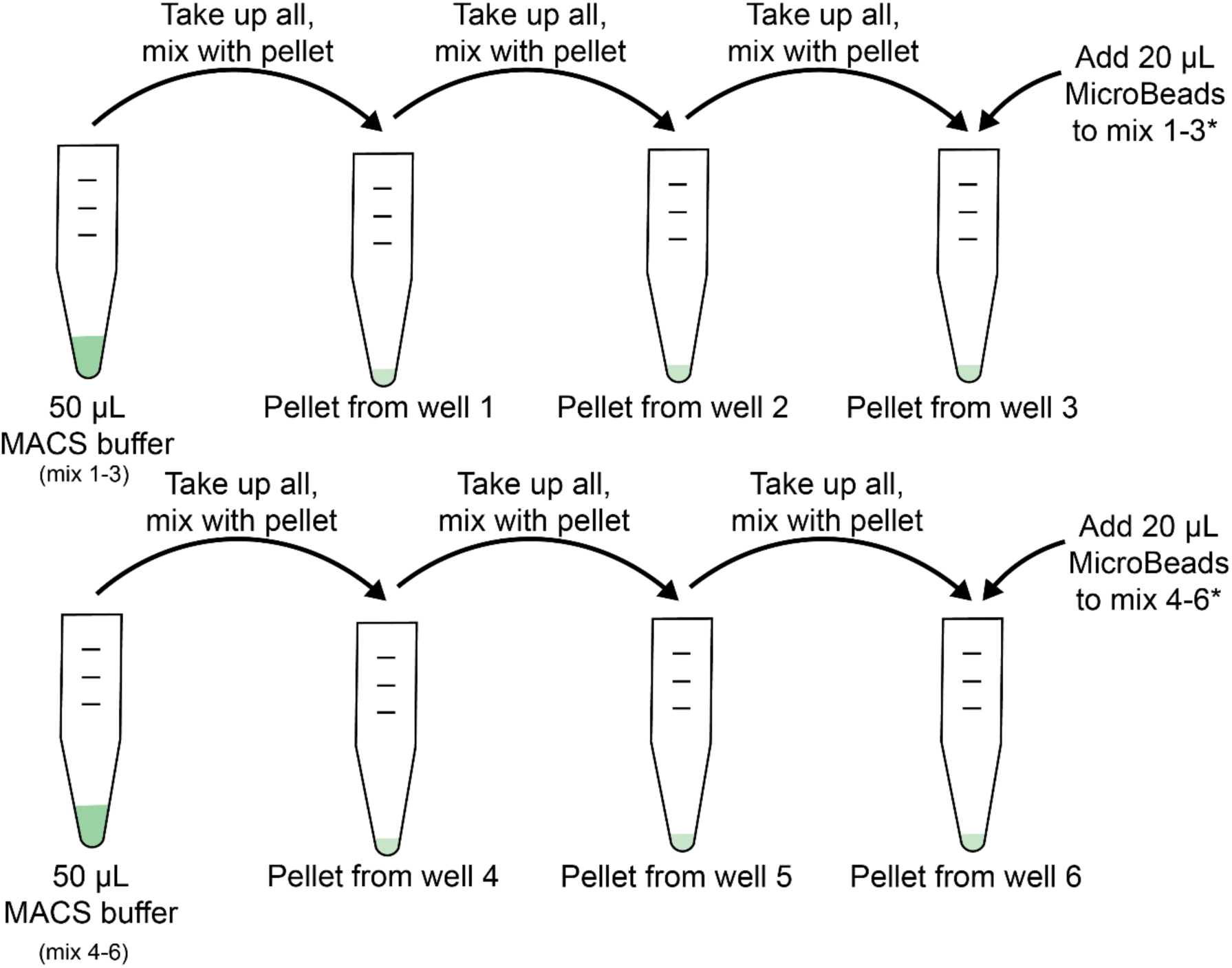
Workflow schematic for combining pellets before the addition of CD144 (VE-cadherin) MicroBeads. The workflow corresponds to processing 6 wells of differentiated endothelial cells, split across two replicates as outlined in *Step 43* and the accompanying *Note*. The asterix (*) relates to the tube layout in Figure 8, specifying the exact tubes used for MicroBeads addition.

**Note**: At this step, the pellets corresponding to the same sample will be combined into one. For example, if processing three 6-wells, the resulting pellets in three individual 1.5 mL Eppendorfs will now be resuspended and combined into the same 50 µL. As we usually process at least six 6-wells, we generate at least two sets of replicate solutions, each resuspended in 50 µL of MACS buffer (see Figure 9 for a workflow schematic). The combined volume of pellets and 50 µL MACS buffer following resuspension should be approximately 100 µL.

44. Take the vial of VE-cadherin MicroBeads out of the fridge and swirl to mix.
45. Add 20 μL of the MicroBeads to each ∼100 µl cell suspension with a P20 pipette, then mix quickly with a P200 pipette by taking the suspension up and down 5 times.
46. Incubate for 15 minutes in the fridge (4°C).
47. Following incubation, add 1.4 mL of cold MACS buffer to the tube with cells and MicroBeads and centrifuge at 200 g for 3 minutes in a swinging-bucket rotor centrifuge for efficient pelleting.

**Note**: If your centrifuge does not have inserts for smaller tubes, use empty 15 mL Falcon tubes as adaptors.

48. Aspirate all supernatant, taking care not to disturb the pellet.
49. Flick the tube gently to resuspend the pellet.
50. Add 500 μL of cold MACS buffer and mix gently to combine.
51. Take up the whole cell solution and pass through a 40 μm strainer and into the original, sample-specific 1.5 mL Eppendorf tube used for the 50 µL MACS buffer aliquots. Keep the cell suspension on a cold block.
52. Put the LS column on a magnet and rinse with 3 mL of cold MACS buffer. Let the solution run through into the collection tube.

**Note**: The number of columns to use per sample will be exactly the same as the number of Eppendorfs prepared with 50 µL MACS buffer in *Step 36c* and, as before, will depend on the initial number of wells used for cell dissociation. For example, for six 6-wells, prepare two LS columns. For details on setting up the MACS kit, visit the manufacturer’s website. For example, use the QuadroMACS Kit by Miltenyi Biotec, with setup photographs available on the website as well.

53. Once the rinse has passed through, apply each 500 µL cell suspension directly onto the LS column and top up with 3 mL of cold MACS buffer. Allow the column to empty into the collection tube fully.
54. Repeat the 3 mL MACS buffer addition 2 more times for a total of 3 washes; allow the column to empty fully between each wash.
55. Carefully remove the column from the separator and place it over a 15 ml Falcon tube.
56. Apply 5 mL MACS buffer and push through using the supplied plunger.

**Optional**: Alternatively, allow the MACS buffer to empty into the new collection tube on its own. This is beneficial when processing multiple samples at once.

57. Take up the cell suspension for counting to determine volume for resuspension for seeding of subconfluent wells.
58. Once counted, centrifuge the cells at 200 g for 3 minutes in a swinging bucket rotor centrifuge for more efficient pelleting.
59. After centrifugation, remove the supernatant and resuspend the cells in the required amount of EGM-2 Complete Medium.
60. Remove the coating from the receiving 12-well plates.
61. Dispense 1 mL of the final cell suspension per well and allow to settle in the hood for 10 minutes before moving to the incubator for a more even cell distribution.

**Note**: We recommend seeding a minimum of 250,000 cells per 12-well (1 mL) and ideally up to 500,000 if possible. The higher cell number will result in confluence upon seeding, which is beneficial for endothelial cell recovery. If seeding 250,000 cells, confluence will be reached within 2 days. Lower seeding numbers are not recommended, especially for ALECs, as these are naturally more quiescent and only proliferate for the first couple of days after MACS. If the cells are seeded too sparsely, they senesce and should not be used for experimentation.

**Critical:** Processing time is extremely important here – the removal of laminin coating needs to be followed with cell addition as soon as possible to prevent drying out. Ensure that the cell suspension is mixed well before dispensing and repeat the mixing of the source solution after dispensing into four wells to avoid settling of the cells in the stripettes.

#### Endothelial cell maturation

**Timing:** 5 days (30 min per day)

Following cell sorting, cells are allowed to undergo a maturation period to stabilise cell identity and allow recovery from sorting-associated stress. This maturation phase is necessary to ensure robust expression of endothelial markers and functional signalling before downstream applications.

62. Following 16–24 h of incubation, refresh the EGM-2 Complete Medium and subject the pure cells to pulsatile (non-oscillatory) flow by using a plate rocker approved for use inside the incubator. We have achieved flow using the RoboRocker 212 (NextAdvance), detailed in *Materials and Equipment*.

**Note**: We apply flow using a RoboRocker 212 programmed to generate continuous pulsatile (but non-oscillatory) motion, with a tilt rate of 5°/second to a maximum angle of 17°, followed by a return to 0° at 1°/second. For a 12-well plate, these parameters correspond to an estimated shear stress of 2 × 10⁻³ Pa. A script used to calculate shear stress under different conditions is available on GitHub (Morison 2025).

**Critical:** We use these rocker settings for ALECs and VLECs cultured in 12-, 24-, and 96-well plates; they are not appropriate for 6-well plates, however, as the resulting medium redistribution will expose cells to drying.

63. Repeat for 4 more days, for a total of 5 days of maturation.

**Note:** We strongly recommend replenishing the medium every day to minimise evaporation; this is crucial under flow as evaporation is accelerated. As with the iPSCs, we recommend consistent daily timings of feeding. After media changes, plates should be returned to the rocker promptly to avoid disrupting flow conditions.

For downstream applications, endothelial cells can be used from 2 days post-MACS onwards, allowing time for recovery, to study endothelial cells at less mature stages.

#### Endothelial cell handling for downstream applications

**Timing:** Variable

This section describes experimental handling and evaluation of the endothelial cells following maturation. Cells generated using this workflow are suitable for downstream functional assays, including in vitro angiogenesis. The steps described here are limited to general handling and splitting of endothelial cells in 12-well plates. Downstream functional assays should be performed using established or independently developed workflows. The tube assay examples shown in Figure 10 were generated using the publicly available Thermo Fisher Scientific workflow (‘Endothelial Cell Tube Formation Assay - UK’).

64. Prepare in advance:

a. The required volume of the EGM-2 Complete Medium appropriate for the endothelial lineage (see *Differentiation media preparation, Step 32 Critical*).
b. A tissue-culture treated plate or vessel of your choosing; check specific downstream workflow for additional coating requirements.
c. EGM-2 Complete Medium and StemPro Accutase pre-warmed to room temperature before starting.

**Figure 10.**
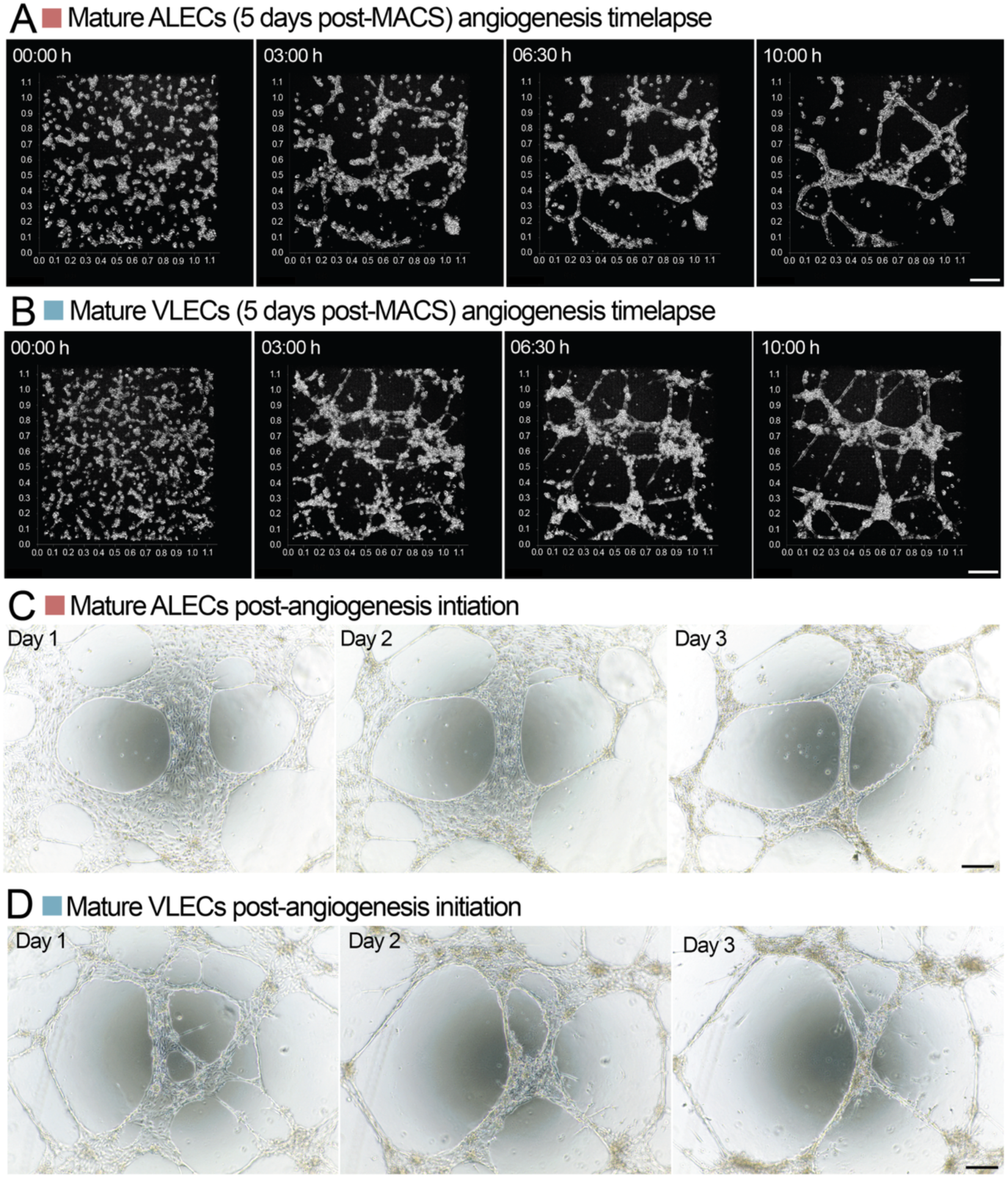
In vitro tube formation assay using iPSC-derived arterial-(ALECs) and venous-like endothelial cells (VLECs). Still images of hourly recordings of mature ALECs (**A**) or VLECs (**B**) seeded at 20,000 cells in a 96-well coated with Geltrex (Geltrex Flex LDEV-Free Reduced Growth Factor Basement Membrane Matrix, Thermo Fisher Scientific Cat# A4000046702). The cells were processed for seeding 5 days post-MACS. Still images were generated from brightfield z-stack confocal live-cell imaging (background subtracted) of the GFP-tagged, endogenous VE-Cadherin, displayed in the XY plane. Daily brightfield micrographs of tube formation from mature ALECs and VLECs are shown in (**C**) and (**D**), respectively. Days refer to days post-seeding for the angiogenesis assay. The data are representative of 2 independent experiments and an additional experiment with 2 independent clones performed using a different matrix (growth factor-reduced Matrigel, Corning Cat# 356231). Scale bar (all images): 200 μm.

**Note**: The required materials and reagents will vary depending on the downstream application and thus are not specified explicitly.

65. Wash each 12-well to be split with 1 mL of DPBS.
66. Treat with 250 μL StemPro Accutase and return to the incubator (37°C) for 3-4 minutes.
67. Return the plate with dissociated cells to the biosafety cabinet, then open and tilt the plate at a 45° angle. Using a P1000 pipette, gently take up the StemPro Accutase solution with cells, then dispense back across the well in a systematic motion that goes from one side to the other, from the top to the bottom of the well, whilst taking care not to pipette over the same area excessively. After doing this 4-5 times, you should observe that most cells have dissociated from the well, and the entire solution can now be collected into a sterile tube pre-filled with 3 times the amount of Accutase.

**Critical**: Do not scratch the surface of the plate. Instead, wash the well gently just above to avoid agitating the cells.

68. Process as required.

**Note**: Endothelial cells generated using this workflow can be difficult to pellet. We therefore recommend centrifugation at 500 g for a minimum of 3 minutes using a swinging-bucket rotor centrifuge with Eppendorf tube adaptors (or open 15 mL Falcons as suitable alternatives).

## Results

This workflow results in arterial-like (ALECs) and venous-like endothelial cells (VLECs), as defined by loss of pluripotency markers and acquisition of lineage-specific markers (Figure 11, **Supplementary Figures 1-2**). As a minimum, we recommend immunofluorescence for VE-cadherin, SOX17 and NR2F2 as the most robust and reproducible readout of differentiation efficiency which also allows for confirmation of correct localisation. By contrast, flow cytometry-based quantification is not localisation-specified and can be confounded by subjective gating strategies and inadvertent exclusion of off-target populations based on cell size, an issue noted in some prior studies where the full gating strategy was made available.

**Figure 11.**
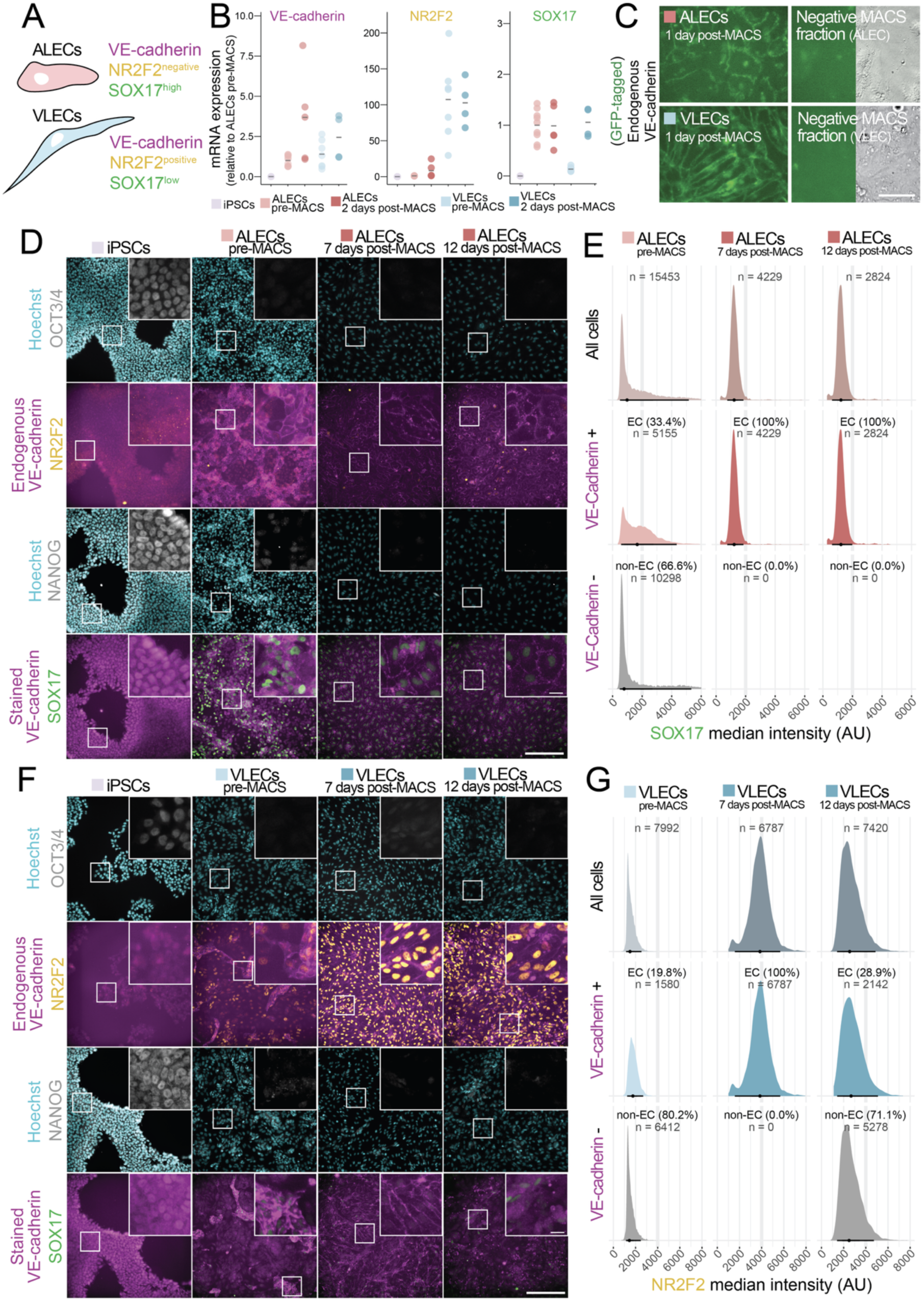
Evaluation of lineage specificity in arterial-(ALECs) and venous-like endothelial cells (VLECs) generated from parental WTC11 GFP-tagged VE-cadherin iPSCs. **A**. Schematic of the expected core lineage marker signature for each endothelial subtypes (Sissaoui et al. 2020; Corada et al. 2013). **B**. RT-qPCR assessment of lineage-specific mRNA marker expression in ALECs and VLECs pre-and post-MACS (magnetic-activated cell sorting). The data are representative of minimum 2 independent iPSC clones and 4 independent experiments, except for ALECs post-MACS (3 experiments) and VLECs post-MACS (1 experiment). **C**. Endogenous GFP-VE-cadherin expression in ALECs, VLECs and the corresponding flow-through fractions (negative control), captured by epifluorescence. Brightfield images are also included for the flow-through to visualise the non-endothelial morphology of the negative cell fraction. Scale bar: 50 μm. **D**. Pluripotency (OCT3/4, NANOG), arterial (SOX17) and venous (NR2F2) marker expression in iPSCs and ALECs across the differentiation time-course, measured using immunofluorescence. Scale bar for insets: 50 μm. Scale bar for main micrographs: 200 μm. **E**. Single-cell quantification of SOX17 nuclear expression in all cells (top) at the indicated ALEC differentiation stage, or specifically in endothelial (VE-cadherin^pos^, middle) versus non-endothelial cells (VE-cadherin^neg^, bottom). The indicated proportion of ECs (33.4%) at the pre-MACS stage was also corroborated by independent cell counting of the subsequent MACS fractions (37.5% VE-cadherin^pos^ cells). **F**. As in (D) in the context of VLEC differentiation. **G**. Single-cell quantification of NR2F2 nuclear expression in all cells (top) at the indicated VLEC differentiation stage, or specifically in endothelial (VE-cadherin^pos^, middle) versus non-endothelial cells (VE-cadherin^neg^, bottom). The indicated proportion of ECs (33.4%) at the pre-MACS stage was also corroborated by independent cell counting of the subsequent MACS fractions (19.4% VE-cadherin^pos^ cells). Histograms in (E) and (F) show single-cell distributions; median and interquartile range (IQR) are indicated by the horizontal bar underneath each histogram. Nuclei were segmented with StarDist (Schmidt et al. 2018), median SOX17 or NR2F2 intensity was measured per nucleus, and nuclei lacking VE-cadherin association were manually excluded to obtain VE-cadherin–positive and –negative cell populations. The data are representative of 3 independent clones and 3 independent replicates. Full staining panels, including additional markers, across independent differentiation time course experiments are included in Supplementary Figures 1 and 2.

With the current implementation, differentiation efficiencies are moderate: 20-60% for VLECs and 30-70% for ALECs prior to magnetic-activated cell sorting (MACS). These values are corroborated by post-MACS cell counts of VE-cadherin-positive and-negative fractions and are reproducible across independent experiments, operators and cell lines, including CRISPR-edited lines harbouring activating *PIK3CA* mutations (not shown). Following MACS, ALECs retain their identity for at least 12 days (Figure 11). VLECs retain their identity for the first week; however, progressive dedifferentiation is observed thereafter, with venous endothelial cells comprising ∼29% of the population by day 12 post-MACS (Figure 12).

**Figure 12.**
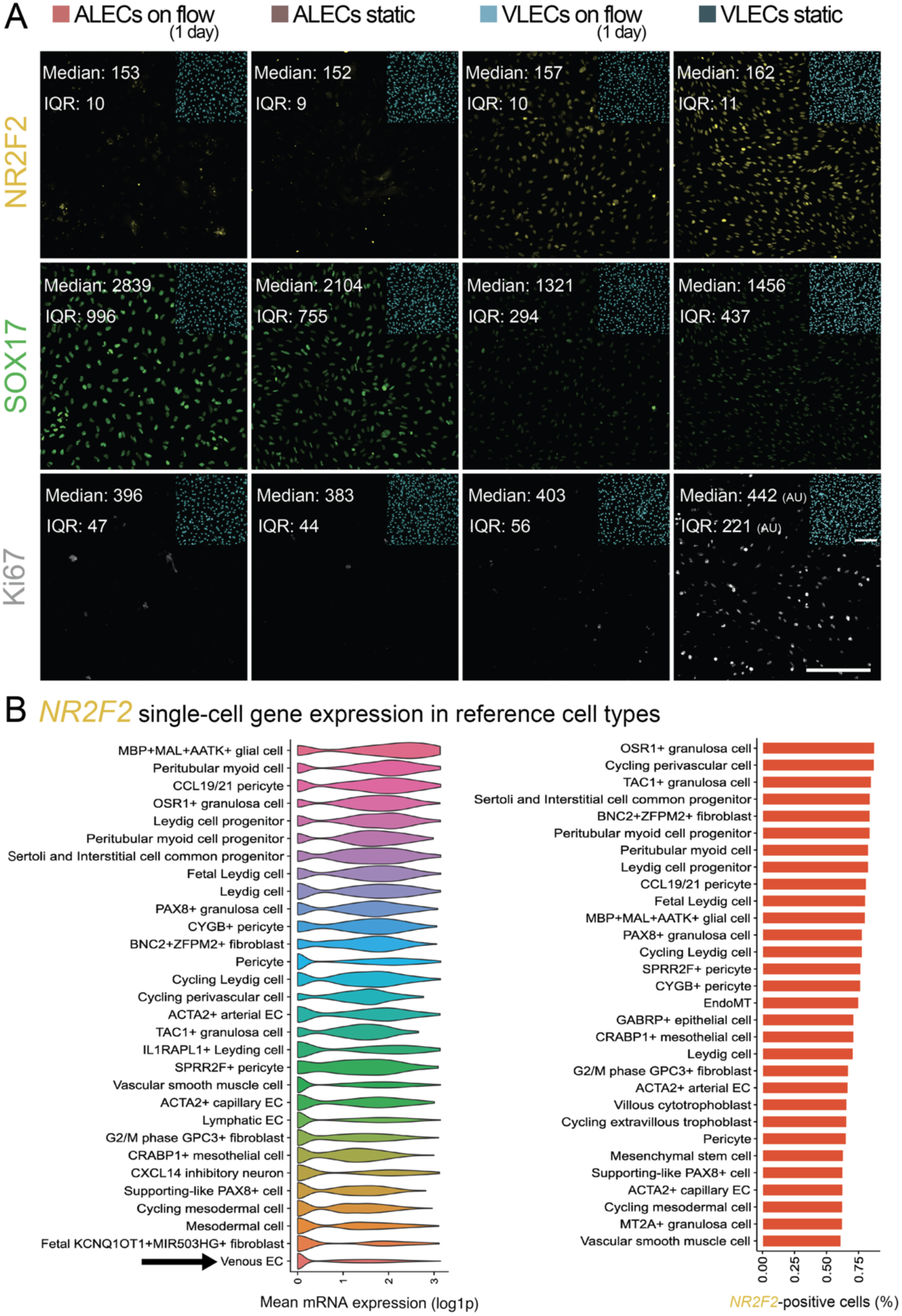
Lineage marker expression in ALECs and VLECs cultured under static versus flow conditions. **A**. All cultures were assessed 2 days post-MACS, with flow exposure for 24 hours before assessment where relevant. ALECs and VLECs were generated from parental WTC11 iPSCs, seeded at 15,000 cells per 96-well (coated with ECMatrix-511 Silk E8 Laminin Substrate). Top panel: expression of the venous NR2F2 marker is low or at background levels in ALECs under both static and flow conditions. In VLECs, NR2F2 expression is higher, with a modest increase observed in static conditions, in line with a concomitant surge in proliferative index as indicated by the extent of Ki67-positive cells (bottom panel). Middle panel: SOX17 expression is enriched in ALECs under flow and reduced under static conditions. Conversely, VLECs show higher SOX17 expression under static culture, indicating emerging loss of lineage fidelity and potential acquisition of an aberrant, mixed arteriovenous phenotype. Insets show Hoechst staining for nuclear identification. Median and interquartile (IQR) staining intensity values for each marker represent quantification of a minimum of 9000 single cells, across 9 independent fields of view. The data are representative of 2 independent clones. Scale bar: 200 μm. **B**. *NR2F2* mRNA expression across annotated human cell types visualised using DISCO (https://www.immunesinglecell.com/genepage/NR2F2), a public resource compiling previously published single-cell RNA-sequencing datasets (Li et al. 2025). Histograms represent the top 30 *NR2F2*-expressing lineages by mean gene expression per cell (log-normalised); barplots show the top 30 *NR2F2*-expressing lineages by percentage-positive cells.

For studies of the developmental consequences of genetic PI3K activation, intermediate differentiation efficiencies are advantageous, as they provide a dynamic range that enables detection of both positive and negative lineage enrichments. For applications requiring higher purity or yield, further optimisation will be required. We are currently developing a three-dimensional version of the workflow to improve differentiation efficiency and enable interrogation of cell-cell interactions in more physiologically-relevant settings.

## Limitations

### Differentiation yield

Differentiation efficiency is sensitive to the quality of the starting iPSC cultures. Reduced pluripotency, diminished viability, or suboptimal culture conditions prior to differentiation initiation will lead to variable and inconsistent outcomes. In addition, we have found that the iPSC culture density before processing for single-cell seeding can influence subsequent proliferation dynamics, confluence and thus overall differentiation efficiency. Differentiation efficiency can also vary as a function of iPSC line, especially if they are engineered to harbour pathological mutations. We have tested the workflow across female and male iPSC lines, wildtype as well as *PIK3CA*-mutant. The specified volumes and yields are based on the average performance of the male WTC11 iPSC line on a *PIK3CA* wildtype genetic background. In line with the original work by Ang et al. (2022), we find that 2D differentiation of VLECs is less efficient than ALECs. We recommend the VE-cadherin MACS step to ensure consistently pure populations of ALECs and VLECs. However, a caveat of MACS is the loss of cells during processing and thus a reduction in the final yield of differentiated cells.

Different culture workflows and coating substrates exist for iPSCs and will impact differentiation efficiency. In this workflow, we opted for commercially available and fully defined reagents, namely StemFlex and ECMatrix-511 Silk E8 Laminin Substrate. We also tested the workflow using iPSCs cultured in Essential 8 Flex Medium (Thermo Fisher Scientific #A2858501) on plates coated with 10 µg/cm^2^ Cultrex Stem Cell Qualified Reduced Growth Factor Basement Membrane (R&D Systems #3434-010-02). We noted consistently poorer survival following single-cell seeding and removal of RevitaCell 24 h later, which compromised the robustness of subsequent endothelial cell differentiation.

### Long-term maintenance and maturation

In general, ALECs and VLECs generated and maintained according to this workflow should not be kept for more than 7 days post-MACS, as they may begin to lose their phenotype, especially if subjected to repeated passaging. We routinely assess lineage stability by immunofluorescence of spare cells seeded in 96-well plates at the time when the main cultures are used for experiments, and recommend that others do the same to ascertain continuous lineage purity. We have consistently noted improved long-term lineage stability when the cells are exposed to pulsatile, albeit gentle (and thus non-physiological) flow on the RoboRocker (**Figure 12A**). Although culture under static conditions leads to a modest increase NR2F2 expression in VLECs, it is also accompanied by a dramatic increase in Ki67-positive nuclei and a partial increase in SOX17 expression, suggesting an aberrant, mixed-lineage identity. Furthermore, NR2F2 is widely used to distinguish venous from arterial endothelial cells, however, it is also highlly expressed in many non-endothelial cell lineages (**Figure 12B**).

It is possible that further improvements in lineage stability and thus long-term culture can be achieved by exposure to physiologically relevant arterial and venous shear stress levels. Nevertheless, one should bear in mind that endothelial cells and ALECs, in particular, are quiscient as expected physiologically, and succesfsul long-term maintenance without artificial immortalisation is unlikely.

The endothelial culture medium used here post-MACS contains 2% fetal bovine serum; alternative, serum-free differentiation strategies have been described (Rosa et al. 2019; Sriram et al. 2015) and could be incorporated in future optimisations to ensure fullly xeno-free culture conditions post-differentiation.

Generally, iPSC-derived endothelial cells present with an immature phenotype, with further maturation taking place upon exposure to flow. In the context of PI3K-driven vascular disorders, the more immature nature of the cells could be advantageous as it may better recapitulate developmental processes.

## Troubleshooting

### Poor differentiation efficiency

Low differentiation efficiency may result from multiple factors, including variability in starting iPSC quality, culture density, or culture conditions before differentiation initiation. While some variability in differentiation efficiency is expected, consistently achieving efficiencies below 20% should prompt troubleshooting. Potential causes may include:

- **Suboptimal quality of the starting iPSC culture**. Initiating experiments with low-quality or partially differentiated cells can compromise downstream results. Maintaining cell health is therefore critical and can be supported by using freshly prepared culture medium, in which supplements are mixed into the base medium shortly prior to use. This approach helps preserve buffering capacity and nutrient stability. In our experience, cells show improved performance when cultured in freshly prepared medium, and aliquoting supplemented medium is an effective strategy to maintain media integrity and consistency over time.
- **Suboptimal seeding density**. Initial seeding density is critical for efficient differentiation. In our experience, cultures seeded too sparsely show poor viability and reduced differentiation efficiency, and the cells generally perform better when plated at higher density (i.e., with sufficient cell–cell contact). This is particularly important for arterial differentiation, where a substantial proportion of cells may be lost during commitment to the arterial lineage; starting with an adequate density helps maintain cell health and supports robust differentiation outcomes. In addition, following MACS, it is important to maximise cell–cell contact; not doing so can lead to rapid senescence and a marked decline in cell quality. This finding aligns with previous reports demonstrating that higher seeding density and increased cell–cell contact promote optimal differentiation and functional properties in iPSC-derived brain microvascular endothelial cells (Wilson et al. 2015).
- **Geometry of the well**. We observe that differentiation efficiency and the extent and timing of cell death can vary between plate formats (e.g., 6-well vs 96-well). This likely reflects format-dependent differences in the local culture microenvironment, including media height (volume-to-surface-area), gas and nutrient exchange, dilution or accumulation of cell-secreted factors, and evaporation or edge effects (particularly in 96-well plates). While we do not systematically optimise for all plate formats, we recommend that users be aware of these effects and consider plate geometry, vessel size, and manufacturer when comparing results or transferring the workflow across different culture formats.
- **Suboptimal quality of differentiation reagents**. Careful tracking of differentiation reagents and their expiry is essential. Reported stability windows may differ depending on formulation, storage conditions, and handling, and users are encouraged to validate reagent stability themselves where feasible. For instance, despite reported instability in water, we have successfully used NKH477 (forskolin analogue) for extended periods at 4°C without detectable degradation, as verified by NMR. Users should independently confirm stability under their own conditions.
- **Temperature or CO_2_ fluctuations**. Differentiation efficiency is highly sensitive to temperature and CO₂ stability. Even small or transient deviations from set conditions can result in pronounced suboptimal cell health and reductions in differentiation efficiency. For example, we have observed dramatic losses in differentiation performance during periods of incubator instability. Users should therefore minimise incubator door openings, limit the time cells spend outside the incubator, and closely monitor incubator temperature, CO₂, and humidity levels throughout the differentiation process.

### Evaluation of VE-cadherin expression

Antibody-based VE-cadherin staining should be interpreted with caution and evaluated in conjunction with cellular morphology and protein localisation, not solely based on the presence of a signal. Correct VE-cadherin localisation is characterised by sharp, continuous appearance at cell-cell junctions, producing well-defined membrane borders, as observed in endothelial cells in Figure 11. In contrast, although signal is also detectable in iPSCs in Figure 11, the staining pattern is diffuse and distributed throughout the cell rather than restricted to the membrane border, reflecting non-specific antibody binding as confirmed by the lack of endogenous signal from the GFP-tagged VE-cadherin allele.

Detection of VE-cadherin may also be influenced by antibody accessibility and epitope availability. We have consistently observed that in arterial-like endothelial cells, the antibody-based VE-cadherin signal is often weaker relative to the endogenous VE-cadherin reporter, whereas in venous endothelial cells, antibody staining can appear comparatively stronger. This likely reflects the tighter intercellular junctions in arterial cells, leading to steric hindrance or reduced epitope accessibility. Notably, we have observed that antibody-based detection of VE-cadherin is more variable across samples, in contrast to detection of the GFP-tagged endogenous protein. Additionally, in cells with weak VE-cadherin expression, the antibody may fail to detect a signal altogether, as observed in **Figure 11F** for VLECs 12 days post-MACS. As a result, tagging endogenous VE-cadherin with a fluorescent reporter provides a more reliable readout of its expression and junctional localisation compared to antibody-based staining alone.

## Resource availability

Further information and requests for resources and reagents should be directed to and will be fulfilled by the lead contact, Dr Ralitsa R. Madsen.

## Acknowledgements

Funding sources: This work was funded by CLOVES Syndrome Community (RRM), the Wellcome Trust (Sir Henry Wellcome Fellowship 220464/Z/20/Z to RRM), UKRI (Future Leaders Fellowship MR/Y017439/1 to RRM) and MRC (PhD studentship to ONM, MRC PPU core funding grant MC_UU_00038).

Collaborators and core facilities: We would like to thank Professor Robert Semple for helpful discussions and for sharing the parental WTC11 iPSC line with us. We thank Miss Sweta Swaminathan for her help with brightfield image acquisition of iPSC maintenance cultures. We thank all members of the Madsen Lab for support with iPSC maintenance, data discussions and final proofreading of the manuscript. We are grateful to Dr Adrien Morison, University of Glasgow, for his help with developing the Python script for shear stress calculations. Lastly, we are indebted to Dundee University’s Imaging Facility (DIF), the National Phenotypic Screening Centre (NPSC), as well as the MRC PPU Lab Management and Technical Teams. Finally, we acknowledge Ang et al. and Loh et al. and their respective teams for establishing the original ALEC/VLEC differentiation protocols, upon which this work builds. Their prior efforts and documentation were essential to the success of the present study.

The authors acknowledge the use of ChatGPT (OpenAI) for assistance with textual revisions. The authors retain full responsibility for the content, accuracy and integrity of the manuscript.

## Author contributions

Conceptualisation: RRM; Methodology: ONM, RRM; Data curation: ONM, RRM; Formal analysis: ONM, RRM; Investigation: ONM, RRM; Project administration: RRM; Resources: RRM; Supervision: RRM; Validation: ONM, RRM; Visualisation: ONM; Writing – original draft: ONM, RRM; Writing – review & editing: ONM, RRM.

## Declaration of interests

RRM has received consulting fees from Nested Therapeutics (Cambridge, U.S.) and serves on the Scientific Advisory Board of CLOVES Syndrome Community.

## Supplementary Figures

**Supplementary Figure 1.**
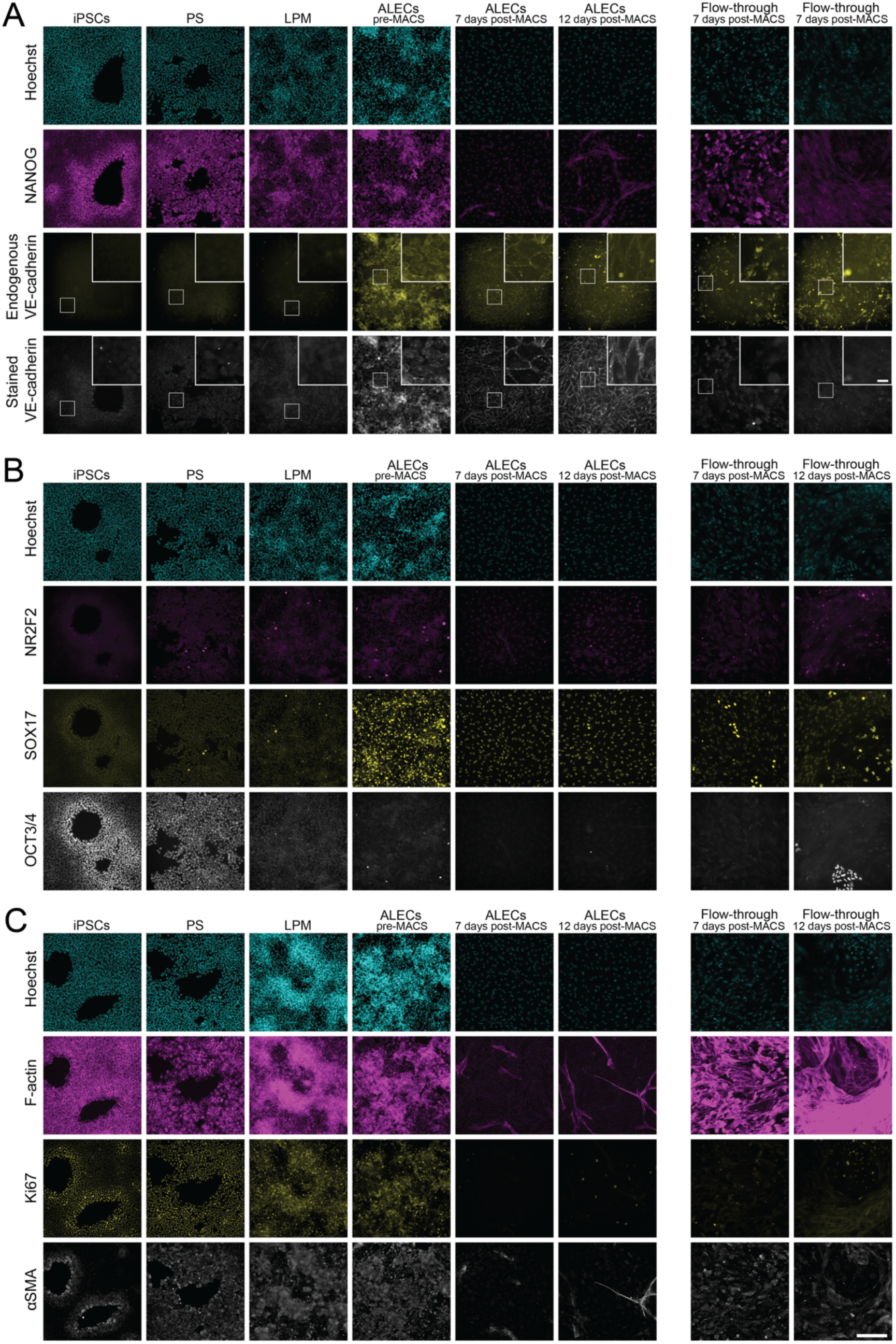
Full staining panel for the differentiation timeline of WTC11 GFP-tagged VE-cadherin iPSCs to mature arterial-like endothelial cells (ALECs). **A**. NANOG (pluripotency marker) and VE-cadherin (endothelial marker) expression at individual stages of iPSC-derived ALEC differentiation, measured by immunofluorescence. VE-cadherin expression in differentiated ALECs is further validated by the fluorescent signal and junctional localisation of the endogenous, GFP-tagged protein. **B**. Immunofluorescence for OCT3/4 (pluripotency marker), NR2F2 (venous marker) and SOX17 (arterial marker in the context of ECs) throughout ALEC differentiation, confirming the expected OCT3/4^neg^NR2F2^neg^SOX17^high^ ALEC signature. **C**. Immunofluorescence for F-actin, Ki67 and αSMA throughout ALEC differentiation. Consistent with their arterial phenotype, pure ALEC cultures present with emerging F-actin-rich tube-like structures and minimal expression for Ki67 and αSMA. All panels also include images from the corresponding MACS flow-through fractions (negative controls), alongside Hoechst staining for cell identification. The data are representative of differentiations from 3 independent clones and 3 experimental replicates, with pure ALECs kept on flow. Scale bar for insets: 50 μm. Scale bar for main micrographs: 200 μm. Abbreviations: ALECs: arterial-like endothelial cells; αSMA: α-smooth muscle actin; iPSCs: induced pluripotent stem cells; LPM: lateral plate mesoderm; MACS: magnetic activated cell sorting; PS: primitive streak.

**Supplementary Figure 2.**
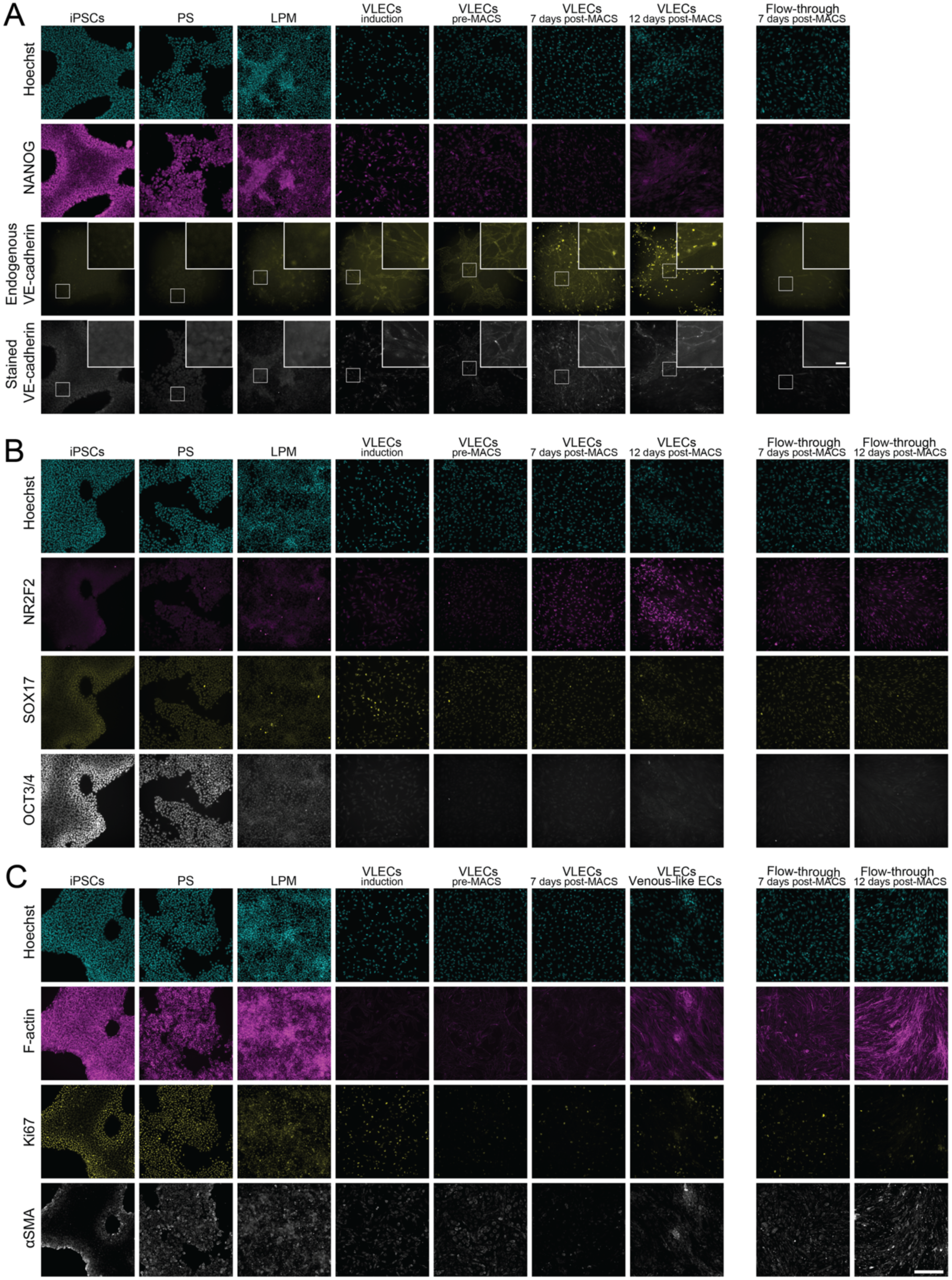
Full staining panel for the differentiation timeline of WTC11 GFP-tagged VE-cadherin iPSCs to mature venous-like endothelial cells (VLECs). **A.** NANOG (pluripotency marker) and VE-cadherin (endothelial marker) expression at individual stages of iPSC-derived VLEC differentiation, measured by immunofluorescence. VE-cadherin expression in differentiated VLECs is further validated by the fluorescent signal and junctional localisation of the endogenous, GFP-tagged protein. **B**. Immunofluorescence for OCT3/4 (pluripotency marker), NR2F2 (venous marker) and SOX17 (arterial marker in the context of ECs) throughout VLEC differentiation, confirming the expected OCT3/4^neg^NR2F2^pos^SOX17^low^ VLEC signature. **C**. Immunofluorescence for F-actin, Ki67 and αSMA throughout ALEC differentiation. Consistent with their arterial phenotype, pure VLEC cultures present with emerging F-actin-rich tube-like structures, moderate expression for Ki67 and minimal αSMA expression. All panels also include images from the corresponding MACS flow-through fractions (negative controls), alongside Hoechst staining for cell identification. The data are representative of differentiations from 3 independent clones and 3 experimental replicates, with pure VLECs kept on flow. Scale bar for insets: 50 μm. Scale bar for main micrographs: 200 μm. Abbreviations: αSMA: α-smooth muscle actin; iPSCs: induced pluripotent stem cells; LPM: lateral plate mesoderm; MACS: magnetic activated cell sorting; PS: primitive streak; VLECs: venous-like endothelial cells.

## Notes

https://www.immunesinglecell.com/genepage/NR2F2

